# A compact spatial map in V2 visual cortex

**DOI:** 10.1101/2021.02.11.430687

**Authors:** Xiaoyang Long, Bin Deng, Jing Cai, Zhe Sage Chen, Sheng-Jia Zhang

## Abstract

Vision plays a critical role in guiding spatial navigation. A traditional view of the visual cortex is to compute a world-centered map of visual space, and visual neurons exhibit diverse tunings to simple or complex visual features. The neural representation of spatio-visual map in the visual cortex is thought to be transformed from spatial modulation signals at the hippocampal-entorhinal system. Although visual thalamic and cortical neurons have been shown to be modulated by spatial signals during navigation, the exact source of spatially modulated neurons within the visual circuit has never been identified, and the neural correlate underpinning a visuospatial or spatio-visual map remains elusive. To search for direct visuospatial and visuodirectional signals, here we record *in vivo* extracellular spiking activity in the secondary visual cortex (V2) from freely foraging rats in a naturalistic environment. We identify that V2 neurons forms a complete spatio-visual map with a wide range of spatial tunings, which resembles the classical spatial map that includes the place, head-direction, border, grid and conjunctive cells reported in the hippocampal-entorhinal network. These spatially tuned V2 neurons display stable responses to external visual cues, and are robust with respect to non- spatial environmental changes. Spatially and directionally tuned V2 neuronal firing persists in darkness, suggesting that this spatio-visual map is not completely dependent on visual inputs. Identification of functionally distinct spatial cell types in visual cortex expands its classical role of information coding beyond a retinotopic map of the eye-centered world.

## Introduction

Vision supports the brain’s allocentric spatial navigation, such as guiding navigation with distal visual cues (Morris, 1981). During navigation, spatial context modulates neuronal responses in the visual cortex (Flossmann and Rochefort, 2020; Nau et al., 2018; Saleem, 2020) and visual thalamus (Hok et al., 2018). Head movement, orientation, direction and locomotion signals were detected in the rodent V1(Bouvier et al., 2020; Guitchounts et al., 2020a; Guitchounts et al., 2020b; Meyer et al., 2018; Saleem et al., 2013; Velez-Fort et al., 2018). Additionally, coherent coding of spatial signals was found in the rodent primary visual cortex (V1) and hippocampal CA1 subfield (Fournier et al., 2020; Haggerty and Ji, 2015; Ji and Wilson, 2007). Vision and movement are known to jointly contribute to hippocampal place code (Chen et al., 2013). Visual cues strongly influence spatial modulation in the hippocampus (Acharya et al., 2016). Place cells, head-direction cells, border cells and grid cells are also all strongly modulated with rotated visual cues (Chen et al., 2019; Hafting et al., 2005; Jeffery and O’Keefe, 1999; Lever et al., 2009; Muller and Kubie, 1987; Solstad et al., 2008; Taube et al., 1990). Meanwhile, all physical location-based spatial cell types have their corresponding visual analogues, which include spatial view cells, saccade direction cells, visual grid and border cells that have been reported in the monkey and human hippocampal-entorhinal network (Doeller et al., 2010; Julian et al., 2018; Killian et al., 2012; Killian et al., 2015; Nau et al., 2018; Rolls and O’Mara, 1995). Spatial tuning has been established in the dorsal lateral geniculate nucleus (dLGN), V1 and other visual cortical areas along the visual pathway from head-fixed as well as freely foraging animals during spatial navigation tasks (Diamanti et al., 2021; Fiser et al., 2016; Fournier et al., 2020; Hok et al., 2018; Jankowski and O’Mara, 2015; Leinweber et al., 2017; Pakan et al., 2018; Saleem et al., 2013; Saleem et al., 2018; Siegle et al., 2021; Taube, 1995; Vantomme et al., 2020). Recent experimental findings have shown that similar hippocampal-entorhinal network mechanisms supporting navigation also mediate a world- centered representation of visual space (Nau et al., 2018). However, a complete function of visual cortical neurons from freely foraging animals in a naturalistic environment remains elusive. We hypothesize that the source of spatio-visual signals is generated from higher-order visual areas in a top-down manner, instead of inheriting directly from the visual thalamus in a bottom-up manner. To test this, we therefore target our recording on V2 instead of the widely studied V1 (Flossmann and Rochefort, 2020; Haggerty and Ji, 2015; Ji and Wilson, 2007). To physiologically characterize the visuospatial responses of visual cortical cells, we performed *in vivo* single-unit recordings in V2, the downstream higher-order structure of V1, while rats freely foraged in a two-dimensional chamber (**Fig. 1a, Supplementary Fig. S1**). Among a total of recorded 1364 well-isolated units, subsets of V2 units showed significant spatial tunings that resemble the place cell, head-direction cell, border cell, grid cell and conjunctive cell in the traditional hippocampal-entorhinal spatial navigation network. The discovery of diverse and compact spatial tunings of V2 neurons provides a more complete spatio-visual map in the visual cortex, suggesting a neural basis of visual spatial coding for spatial navigation.

**Fig. 1.**
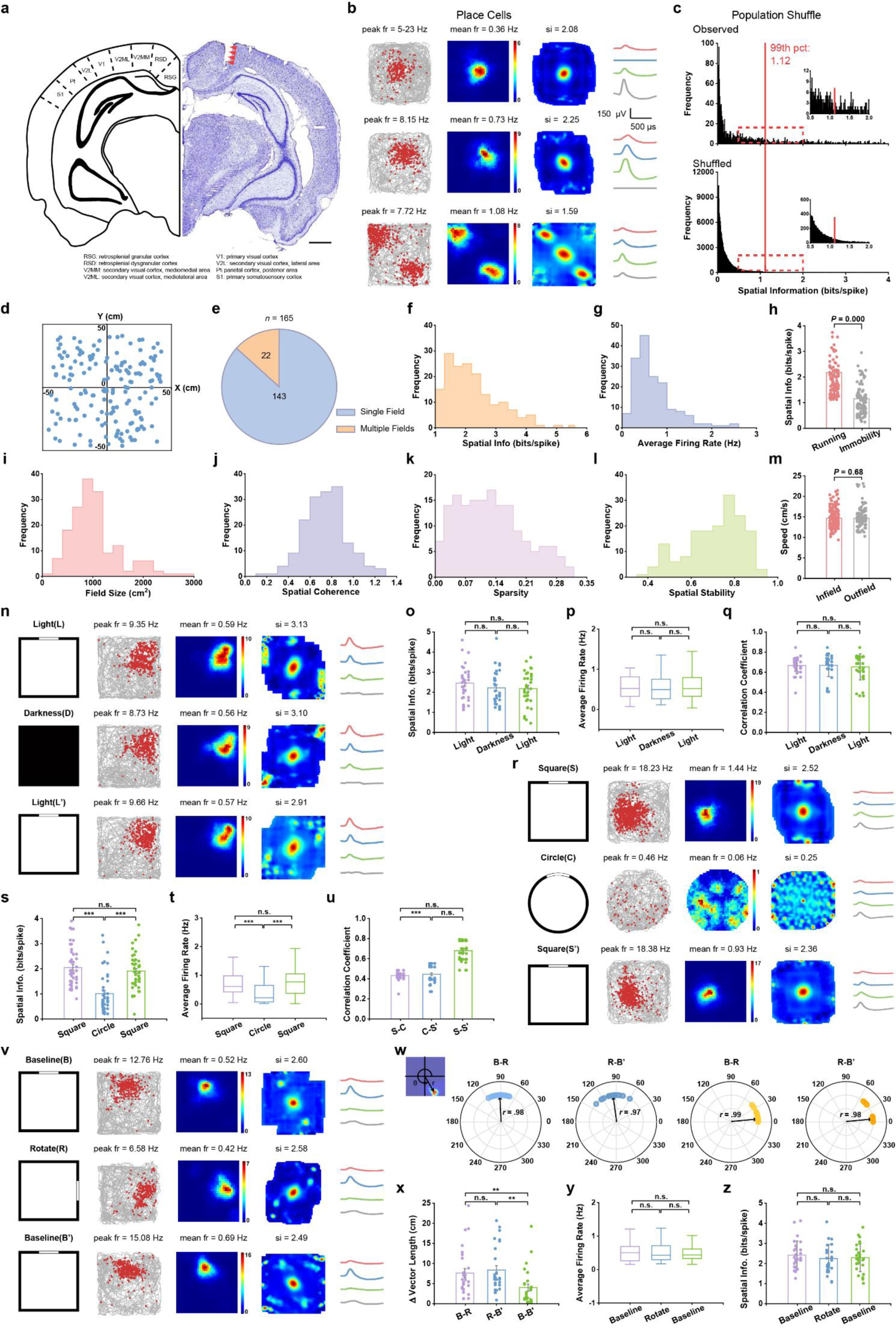
Place cells in the visual cortex. **a**, Photograph of a coronal brain section (right) stained with cresyl violet illustrating the track (arrowheads) of the recording electrodes passing through the rat secondary visual cortex (V2). Schematic diagram (left) around V2. Scale bar, 1 mm. **b**, Three representative V2 place cells. First column: rat’s run trajectory (grey line) with superimposed spike locations (red dots). Second column: spatial firing rate maps. Firing rate is color-coded with dark blue (red) indicating minimal (maximum) firing rate. The two numbers in the middle panel represent the peak and mean firin g rates, respectively. Spatial information (SI) score is labeled at the top of firing rate map. Third column: autocorrelation diagrams. The scale of the autocorrelation maps is twice that of the spatial firing rate maps. Fourth column: average spike waveforms on four electrodes of the same tetrode are shown on the right column. **c**, Distribution of SI for observed data (top panel) and shuffled data (bottom panel). Red line indicates the 99^th^ percentile for the SI score derived from the shuffled data. The inset shows the zoomed-in of the red dashed box. **d**, Distribution of the center of firing fields from all recorded V2 place cells showed uniformly distributed patterns (*n* = 165, Friedman-Rafsky’s MST test, *P* = 0.34, two-tailed *z*-test). **e**, Venn diagram showing the number of V2 place cells with one or more firing fields. **f-g**, Distribution of mean firing rate and SI derived from all V2 place cells. **h**, SI was significantly higher during running than during slow movement or immobility. **i**, Distribution of place field sizes. **j-l**, Distribution of spatial coherence, sparsity and intra-trial spatial stability between the first and second halves of all identified V2 place cells (*n* = 165). **m**, No change in running speed between the within- and outside-firing fields (*n* = 165, two-tailed paired *t*-test, *P* = 0.68). **n**, Firing properties of one representative V2 place cell in the light condition and total darkness. First column: schematic of the experimental paradigms; responses of the place cell across light (upper panel), darkness (middle panel) and back to light condition (lower panel). **o-p**, Both SI score (L-D: *P* = 0.08, D-L’: *P* = 0.97 and L-L’: *P* = 0.11) and mean firing rate (*n* = 32, two-sided Wilcoxon signed rank test, L-D: *P* = 0.60, D-L’: *P* = 0.47 and L-L’: *P* = 0.45) remained stable in the light condition, in total darkness and back to light condition. **q**, Place cells exhibited stable tuning between the switch from light (L) to darkness (D) and then light (L’) conditions (*n* = 32, two-sided Wilcoxon signed rank test, L-D: *P* = 0.50, D-L’: 0.14 and L-L’: 0.90). **r**, Responses of one representative V2 place cell in square enclosure and circle. First column: schematic of the experimental paradigms; firing patterns of the place cell in the square enclosure (upper panels), circle (middle panels) and back to the square enclosure (lower panels). **s-t**, Quantification of SI in the remapping experiment (*n* = 45, two-sided Wilcoxon signed rank test, S-C: *P* = 0.000, C-S’: *P* = 0.000 and S-S’: *P* = 0.74) and mean firing rate (S-C: *P* = 0.000, C-S’: *P* = 0.000 and S-S’: *P* = 0.30). **u**, Spatial correlation showed significant remapping in different environmental shapes (S-C: *P* = 0.000, C-S’: *P* = 0.000 and S-S’: *P* = 0.39). **v**, Responses of one representative V2 place cell under visual cue manipulation. First column: schematic of the experimental paradigms; firing properties of the place cell in the baseline condition (upper panels), counterclockwise 90°cue-rotation manipulation (middle panels) and back to the original baseline condition (lower panels). **w**, Firing properties of place cells following cue card rotation. Upper left panel: the angle and length of the coordinate vector of the field center relative to the center of the box. First and second columns: polar plots showing angular shifts in the direction of the coordinates of the firing field center relative to the box center (*n* = 27, B-R, Rayleigh test, *P* = 0.000, *r* = 0.98, median shift = 92.83°; R-B’, Rayleigh test, *P* = 0.000, *r* = 0.97, median shift = 97.83°; two-sided Wilcoxon signed rank test, *P* = 0.000 and 0.000 for B-R and R-B’, respectively). Third and fourth columns: similar to the first and second columns, except for place cells not following the cue card manipulation. Polar plot showing angular shifts in direction of the coordinates of the firing field center relative to the box center (*n* = 24, B-R, Rayleigh test, *P* = 0.000, *r* = 0.99, median shift = 5.22°; R-B’, Rayleigh test, *P* = 0.000, *r* = 0.98, median shift = 5.37°; two-sided Wilcoxon signed rank test, *P* = 0.95 and 0.42 for B-R and R-B’, respectively). **x**, The difference in vector length was smaller between two baseline conditions (*n* = 27, two-sided Wilcoxon signed rank test, *P* = 0.18, 0.002 and 0.008, respectively). **y-z**, Among baseline, cue card rotation and baseline, neither mean firing rates (*n* = 27, two-sided Wilcoxon signed rank test, *P* = 0.92, 0.51 and 0.73, respectively) nor SI scores (*n* = 27, two-sided Wilcoxon signed rank test, *P* = 0.26, 0.97 and 0.21, respectively) were significantly different.

## Results

We first implanted movable micro-drives connected with four tetrodes into six male Long-Evans adult rats. Recording electrodes were targeted to the higher-order secondary visual cortex. We then recorded spiking activity while rats foraged for randomly scattered ground biscuits in an open two-dimensional square or cylinder enclosure. A total of 1364 well-isolated single units were recorded mainly from deep layers IV to VI (starting around 600 μm below the dura) of V2 across 278 recording sessions. The quality of multiple well-separated clusters and spikes (waveforms) was assessed with two well-established criteria (**Supplementary Fig. S2**). Postmortem histological reconstruction verified that recording electrode positions of all implanted animals were located in V2, with a narrow region approximately 4.5 mm posterior to bregma and 2.5 mm lateral to midline (**Fig. 1a, Supplementary Fig. S1**). The distribution of distinct spatially tuned cell types between V2 cortical layers showed a tendency for place cells, head-direction cells and border cells to cluster in layer-V/VI, while grid cells appeared to distribute evenly in deep layers (**Supplementary Fig. S3**). The number of each functionally distinct V2 spatially tuned cells from individually implanted rats was summarized in **Supplementary Fig. S4**.

### Place cells in V2

A large number of well-isolated V2 units displayed spatially selective firing patterns with respect to the rat’s location in the open field, which was termed as the cell’s “firing field” (O’Keefe and Dostrovsky, 1971). We used a strict criterion to define spatially tuned units. If the statistical criterion was relaxed, a higher percentage of V2 units would meet the threshold criterion. V2 place cells were classified if their spatial information (SI) scores were above a stringent 99^th^ percentile of shuffled rate maps. We identified 165/1364 (12.1%) significant spatially modulated V2 units as “place cells” based on their SI scores (**Fig. 1b, c, Supplementary Fig. S7**) using population shuffling, which is more stringent than within-cell shuffling (**Supplementary Fig. S6**). This percentage was significantly higher than expected by random selection from the entire shuffled population (**Fig. 1c**; Z = 41.19, *P* < 0.001; binomial test with expected *P*0 of 0.01 among large samples). Furthermore, we validated spatial tuning of V2 place cells using a maximum-likelihood approach (**Supplementary Fig. S8**). Together, these V2 place receptive fields uniformly covered the environment (**Fig. 1d**, Friedman-Rafsky’s MST test, *P* = 0.34, two-tailed *z*-test) (Smith and Jain, 1984). The statistics of peak-to-trough spike width and peak-to-peak amplitude of V2 place cells were shown in Supplementary Fig. S5a. A large percentage (143/165, 86.67%) of V2 place cells had a single place field (**Fig. 1e**), where the remaining (26/165, 14.33%) V2 place cells displayed multiple fields (**Fig. 1e**). V2 place cells had 2.17 ± 0.04 bits per spike (all data are shown as mean ±s.e.m. unless specified otherwise) in SI score (**Fig. 1f**). The mean and peak firing rates of V2 place cells were (0.73 ±0.04 Hz) and (9.68 ±0.36 Hz), respectively (**Fig. 1g**). The SI score was significantly greater during running than inactive mobility (**Fig. 1h**, Supplementary S9a, *n* = 165, *P* = 0.000, two-tailed paired *t*-test). The average receptive field size of V2 place cells was 31.98% about the size of the open field (1022.40 ±39.08 cm^2^, **Fig. 1i**). Additionally, V2 place cells had an average spatial coherence of 0.75 ±0.02 (**Fig. 1j**), and an average spatial sparsity of 0.12 ±0.01 (**Fig. 1k**).

Between the first and the second halves of single intra-session recordings, V2 place cells displayed a high degree of spatial correlation (**Fig. 1l**, Supplementary Figs. S10a-c). There was no significant difference between the “in-field” and “out-field” running speed (14.66 ± 0.18 cm/s vs 14.65 ± 0.16 cm/s, *n* = 165, *P* = 0.68, two-tailed paired *t*-test, **Fig. 1m**). Notably, V2 place fields remained stable during three consecutive 60-min light (L)-dark (D)-light (L’) recording sessions (each session for 20 min) (**Fig. 1n** and Supplementary S11). There were no significant changes in SI (**Fig. 1o**, *n* = 32, two-sided Wilcoxon signed rank test, L-D: *P* = 0.08, D-L’: *P* = 0.97 and L-L’: *P* = 0.11), mean firing rate (**Fig. 1p**, same test, L-D: *P* = 0.60, D-L’: *P* = 0.47 and L-L’: *P* = 0.45) and spatial correlation coefficient (**Fig. 1q**, L-D: *P* = 0.50, D-L’: *P* =0.14 and L-L’: *P* =0.90), suggesting that V2 spatial firing had little dependence on the purely visual input. Notably, simultaneously recorded place cells, border cells, grid cells and head-direction cells preserved their spatial firing properties in total darkness (**Supplementary Fig. S12**). In remapping experiments of square (S)-circle (C)-square (S’), we found that V2 place cells remapped with the changed SI (**Fig. 1s** and **Supplementary Fig. S15**, *n* = 45, two-sided Wilcoxon signed rank test, S-C: *P* = 0.000, C-S’: *P* = 0.000 and S-S’: *P* = 0.74), mean firing rate (**Fig. 1t**, same test, S-C: *P* = 0.000, C-S’: *P* = 0.000 and S-S’: *P* = 0.30) and spatial correlation coefficient (**Fig. 1u**, S-C: *P* = 0.000, C-S’: *P* = 0.000 and S-S’: *P* = 0.39), suggesting that the environmental shape influenced the firing properties of V2 place cells. Furthermore, V2 place cells rotated and remained stable during baseline (B)-rotation (R)-baseline (B’) when visual cues were rotated by 90° in orientation (**Fig. 1v** and **Supplementary Fig. S13**), as measured by the angle and length of the coordinate vector of the firing field relative to the center of the arena (*n* = 27, Rayleigh test: for B-R, *P* = 0.000, *r* = 0.98, median shift = 92.83°; for R-B’, *P* = 0.000, *r* = 0.97, median shift = 97.83°; two-sided Wilcoxon signed rank test, *P* = 0.000 and 0.000 for B-R and R-B’, respectively; see the second and third columns in **Fig. 1w** and **Fig. 1x**). However, for those place cells that did not follow the cue rotation, there were very small angular shifts (*n* = 24, Rayleigh test: for B-R, *P* = 0.000, *r* = 0.99, median shift = 5.22°; for R-B’, *P* = 0.000, *r* = 0.98, median shift = 5.37°; two-sided Wilcoxon signed rank test, *P* = 0.95 and 0.42 for B-R and R-B’, respectively; see the third and fourth columns in **Fig. 1w**). There was no significant difference in mean firing rate (**Fig. 1y**, *n* = 27, *P* = 0.92, 0.51 and 0.73 for R-B’, B-R and R-B’, respectively) or spatial information (**Fig. 1z**, *P* = 0.26, 0.97 and 0.21, respectively) from baseline to cue card rotation and then back to baseline. Together with V2 place cells, simultaneously recorded V2 head-direction cells, V2 border cells and V2 grid cells rotated concurrently with the cue card (**Supplementary Fig. S13**). Additionally, simultaneously recorded place cells, border cells, grid cells and head-direction cells on the same microdrive rotated coherently to the visual landmark (**Supplementary Fig. S14**).Therefore, place cells in V2 shared similar spatially selective firing features as the classical hippocampal place cells (O’Keefe and Dostrovsky, 1971).

### Border cells in V2

Border cells fire specifically whenever an animal forages close to one or multiple physically-defined local boundaries of an open field (Lever et al., 2009; Solstad et al., 2008). Next, we examined whether traditional border cells existed in the V2. We assigned a V2 unit as a border cell if its border score (BS) was larger than the 99^th^ percentile of shuffled distribution (**Fig. 2b**). The histograms of peak-to-trough spike width and peak-to-peak amplitude were shown in Supplementary Fig. S5b. Overall, 85/1364 (6.23%) recorded V2 units were categorized as border cells (**Fig. 2a** and **Supplementary Fig. S16**), and this percentage was significantly higher than expected by random selection from the entire shuffled population (**Fig. 2b**; Z = 19.41, *P* < 0.001; binomial test with expected *P*0 of 0.01 among large samples). The BS was higher during animal’s running (> 2.5 cm/s) than immobility (2.5 cm/s) (**Fig. 2d** and Supplementary Fig. S9b, *n* = 85, *P* = 0.000, two-tailed paired *t*-test). V2 border cell had a mean firing rate of 1.51 ± 0.21 Hz (**Fig. 2e**) and an average BS of 0.69 ± 0.01 (**Fig. 2f**). Similar to the classical border cells reported in the medial entorhinal cortex (mEC) and subiculum (Solstad et al., 2008), V2 border cells fired along a single border (*n* = 39/85, 45.88%) while others discharged actively along with two (*n* = 15/85, 17.65%), three (*n* = 3/85, 3.53%) or even four (*n* = 28/85, 32.94%) borders within the open arena (**Fig. 3c** and **Supplementary Fig. S16**).

**Fig. 2.**
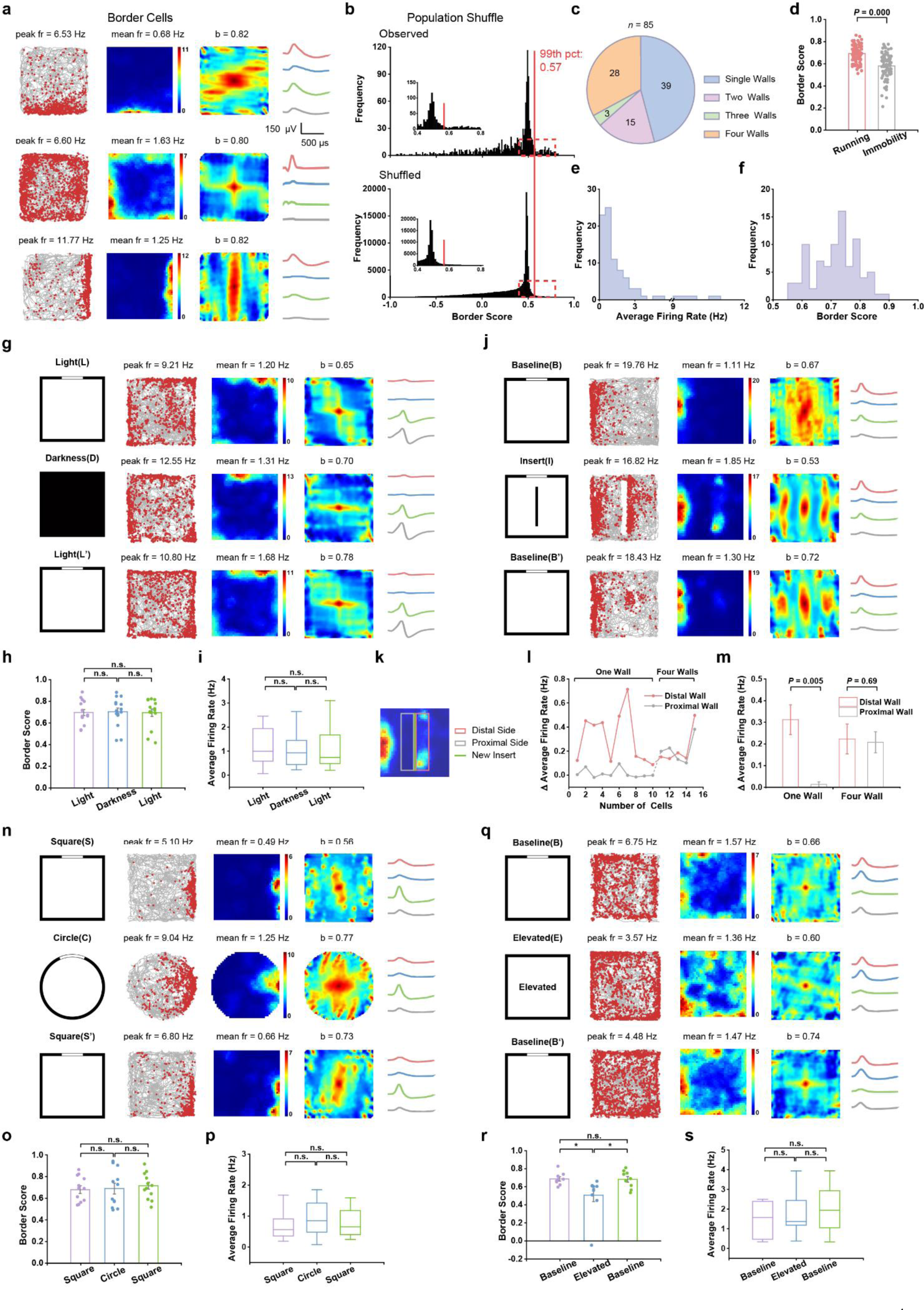
Border cells in the visual cortex. **a**, Three representative border cells identified from the secondary visual cortex. Notations and symbols are the same as those in Fig. 1a. Border score (BS) for each border cell is labelled at the top of firing rate map. **b**, Distribution of BS for observed data (top panel) and shuffled data (bottom panel). Red line indicates the 99^th^ percentile for BS derived from the shuffled data. The inset shows the zoomed-in view of the red dashed box. **c**, Venn diagram showing the number of border cells having firing fields along one, two, three and four walls, respectively. **d**, BS was statistically higher during running than slow movement or immobility (*n* = 85, two-tailed paired *t*-test, *P* = 0.000). **e- f**, Distribution of mean firing rate and BS derived from all identified border cells. **g**, Firing properties of one representative border cell in the light condition and total darkness. Notations and symbols are the same as those in Fig. 1n. **h-i**, Quantification of BS in the L-D-L’ conditions (*n* = 15, two-sided Wilcoxon signed rank test, L-D: *P* = 0.53, D-L’: *P* = 0.69 and L-L’: *P* = 0.73, respectively) and mean firing rate (*n* = 15, two-sided Wilcoxon signed rank test, L-D: *P* = 0.12, D-L’: *P* = 0.31 and L-L’: *P* = 0.59). **j**, Responses of one representative border cell before and after insertion of a new wall. First column: Schematic of the experimental paradigms. Third column: Responses of the border cell from baseline (upper panels), wall insertion (middle panels) and baseline conditions (lower panels). **k**, Schematic showing the method of estimating responses to the insertion of a discrete wall (green line). Firing rates were calculated in both 15-cm-wide distal side (red box) and proximal side (gray box) of the new insert. Proximal and distal sides were determined by the location relative to that of the original border firing field. **l**, Difference in mean firing rate between (distal and proximal) wall insertion and baseline. Data from ten V2 units with firing fields along a single wall and five V2 units with firing fields along four walls. **m**, The distal side had a higher mean firing rate than the proximal side for border cells with firing field along a single wall (*n* = 10, two-sided Wilcoxon signed rank test, *P* = 0.005). For border cells firing along four walls, distal and proximal sides had similar mean firing rates (*n* = 5, two-sided Wilcoxon signed rank test, *P* = 0.67). **n**, Responses of one representative V2 border cell in square enclosure and circle. Notations and symbols are the same as those in Fig. 1r. **o-p**, Quantification of BS (*n* = 12, two-sided Wilcoxon signed rank test, S-C: *P* = 0.94, C-S’: *P* = 0.64 and S-S’: *P* = 0.72) and mean firing rate (S-C: *P* = 0.31, C-S’: *P* = 0.54 and S-S’: *P* = 0.81) in the square enclosure (top panels), circle (middle panels) and back to the square enclosure (bottom panels). **q**, Responses of one representative V2 border cell in baseline (top panels), elevated platform without walls (middle panels) and baseline condition (bottom panels). First column: schematic of the experimental paradigms. **r-s**, Quantification of mean firing rate (*n* = 9, two-sided Wilcoxon signed rank test, S-C: *P* = 0.52, C-S’: *P* = 0.21 and S-S’: *P* = 0.52) and BS during baseline, elevated platform without walls and back to baseline sessions (*n* = 9, two- sided Wilcoxon signed rank test, B-E: *P* = 0.02, E-B’: *P* = 0.05 and B-B’: *P* = 0.86).

**Fig 3.**
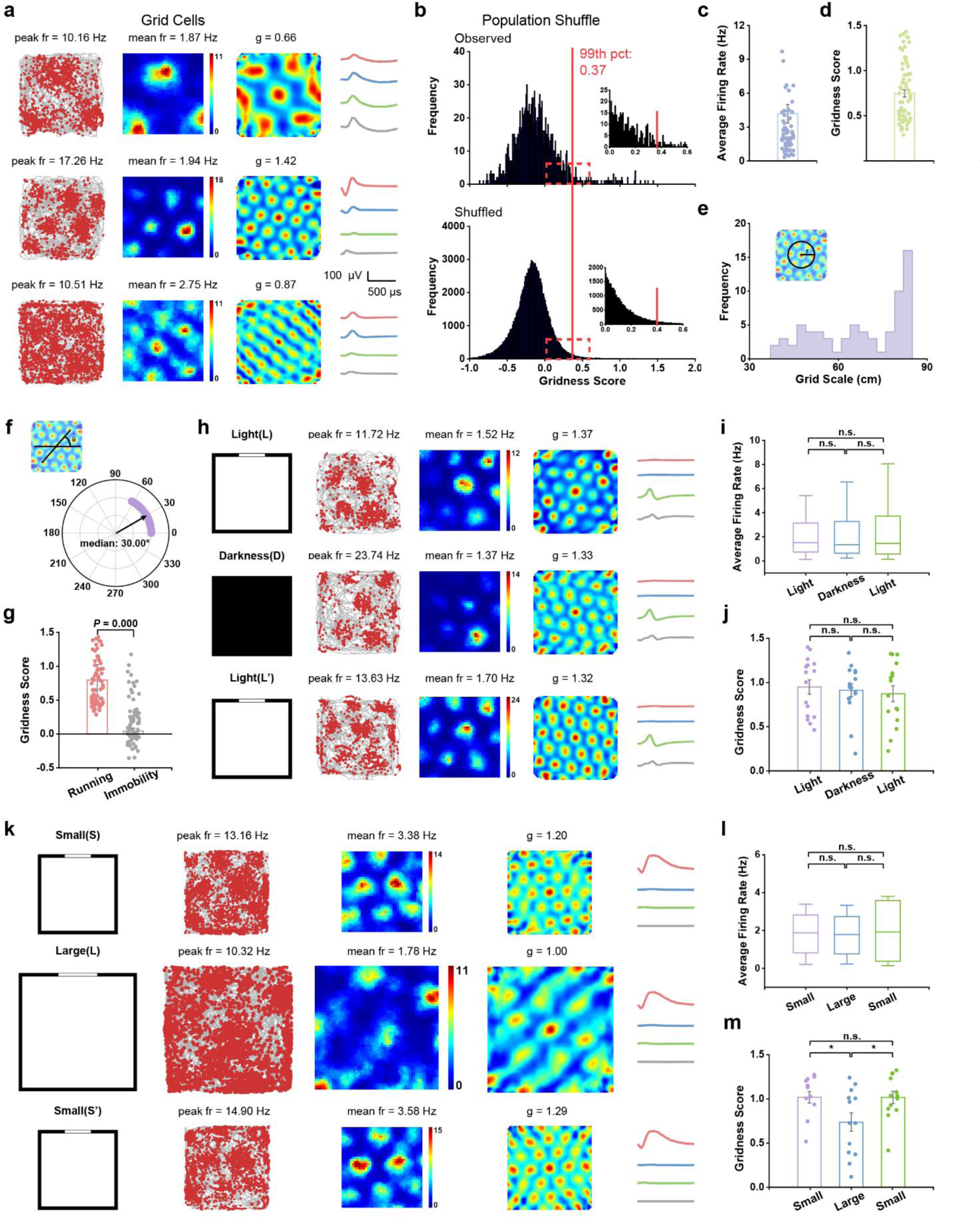
Grid cells in the visual cortex. a, Three representative grid cells recorded from the visual cortex. Similar symbol notations as Fig. la. Gridness score (GS) for each grid cell is labelled at the top of firing rate map. b, Distribution of for observed data (top panel) and shuffled data (bottom panel). Red line indicates the 99th percentile for GS in the shuffled data. The inset shows the zoomed-in view of the red dashed box. **c-d**, Mean firing rate and GS statistics for all identified V2 grid cells. **e-f**, Distribution of gridness scales and grid orientations derived from all identified V2 grid cells. **g**, Grid firing patterns were more pronounced during running than immobility (*n* = 70, two-tailed paired *t*-test, *P* = 0.000). **h**, Firing properties of one representative V2 gird cell in the light condition and total darkness. **i-j**, Quantification of mean firing rate (*n* = 16, two-sided Wilcoxon signed rank test, L-D: *P* = 0.50, D-L’: *P* = 1.00 and L-L’: *P* = 0.80, respectively) and GS in the L-D-L’ conditions (*n* = 16, two-sided Wilcoxon signed rank test, L-D: *P* = 0.57, D-L’: *P* = 0.80 and L-L’: *P* = 0.28, respectively). **k**, Firing properties of one representative V2 gird cell in an enclosure of different size. **l-m**, Quantification of mean firing rate (*n* = 13, two-sided Wilcoxon signed rank test, S-L: *P* = 0.97, L-S’: *P* = 0.92 and S-S’: *P* = 0.97, respectively) and GS in varying sizes of running boxes: 1 × 1 m^2^ (S), 1.5 ×1.5 m^2^ (L) and 1 × 1 m^2^ (S’) (*n* = 13, two-sided Wilcoxon signed rank test, S-L: *P* = 0.023, L-S’: *P* = 0.023 and S-S’: *P* = 0.86). The GS decrease in a larger environment was consistent with the notion that grid cells anchor locally.

During consecutive 60-min light-dark-light sessions (**Fig. 2g** and **Supplementary Fig. S17**), there was no statistically significant difference in BS (**Fig. 2h**, *n* = 15, two-sided Wilcoxon signed rank test, L-D: *P* = 0.53, D-L’: *P* = 0.69 and L-L’: *P* = 0.73) and mean firing rate (**Fig. 2i**, L-D: *P* = 0.12, D-L’: *P* = 0.31 and L-L’: *P* = 0.59), indicating that border tuning in V2 is independent on the visual input. Introduction of a new wall into the square arena produced a new border field on the distal side of the new wall for one-sided border cells (**Fig. 2j**) while the newly inserted wall evoked new border fields on both proximal and distal sides of the wall insert for four-sided border cells (**Figs. 2k-m** and **Supplementary Fig. S20**, *n* = 15) (Solstad et al., 2008). For the same V2 border cell, there was no significant change in mean firing rate (*n* = 12, two-sided Wilcoxon signed rank test, S-C: *P* = 0.31, C-S’: *P* = 0.54 and S-S’: *P* = 0.81) and BS (S-C: *P* = 0.94, C-S’: *P* = 0.64 and S-S’: *P* = 0.72) while navigating between circular and square environments (**Figs. 2n-p** and **Supplementary Fig. S18**). Additionally, removal of four outer walls from the enclosure on the elevated platform preserved similar V2 border fields (**Fig. 2q** and **Supplementary Fig. S19**), suggesting that V2 border cells anchored only to geometric boundaries instead of physical edges. No significant change in mean firing rate was observed between the standard walled enclosure and the elevated wall-free platform (**Fig. 2s**, *n* = 9, two-sided Wilcoxon signed rank test, B-E: *P* = 0.52, E-B’: *P* = 0.21 and B-B’: *P* = 0.52), yet there was a decrease in BS on the elevated platform (**Fig. 2r**, B-E: *P* = 0.02, E-B’: *P* = 0.05 and B-B’: *P* = 0.86). Therefore, V2 border cells resembled those classical border cells reported in the rat mEC and subiculum (Lever et al., 2009; Solstad et al., 2008).

### Grid cells in V2

Hexagonal firing patterns of classical grid cells have been reported in the rat mEC (Hafting et al.,2005). To identify potential grid cells in V2 (**Fig. 3a** and **Supplementary Fig. S21**), we calculated the gridness score (GS, a metric measuring the degree and symmetry of spatial periodicity of hexagonal firing patterns) based on rotated spatial autocorrelation of the firing rate maps. We classified a V2 unit as a grid cell if its GS was higher than the 99^th^ percentile value of the shuffled data. A small percentage (70/1364 or 5.13%) V2 units met this significance criterion (**Fig. 3b**). This percentage was significantly higher than expected by random selection from the entire shuffled population (**Fig. 3b**; *Z* = 15.34, *P* < 0.001; binomial test with expected *P*0 of 0.01 among large samples). V2 grid cells had a mean firing rate of 3.75 ±0.72 Hz (**Fig. 3c**) and an average GS of 0.75 ±0.04 (**Fig. 3d**). We further quantified the properties of V2 grid cells (average grid spacing: 67.44 ±1.28 cm, **Fig. 3e**; median grid orientation: 30.00°, **Fig. 3f**). Additionally, the fluctuation in V2 grid spacing was not a result of interindividual differences between rats (Supplementary **Fig. S4c**).

Furthermore, we found that V2 grid cells showed higher GSs during running (> 2.5 cm/s) than slow mobility (< 2.5 cm/s) (**Fig. 3g** and Supplementary Fig. S9c, *n* = 70, *P* = 0.000, two-tailed paired *t*-test). Additionally, V2 grid cells maintained hexagonal firing patterns in total darkness (**Fig. 3h** and **Supplementary Fig. S22**), and there was no significant change in mean firing rate (**Fig. 3i**, *n* = 16, two-sided Wilcoxon signed rank test, L-D: *P* = 0.50, D-L’: *P* = 1.00 and L-L’: *P* = 0.80) and GS (**Fig. 3j**, L-D: *P* = 0.57, D-L’: *P* = 0.80 and L-L’: *P* = 0.28) during consecutive light-dark-light sessions, suggesting that visual inputs are not essential for maintaining visual grid firing patterns. Discharging patterns of V2 grid cells in a larger environment (1.5 × 1.5 m^2^ box) preserved multiple regular triangular structures and high GS similar to those in a smaller (1 × 1 m^2^ box) enclosure (Supplementary **Figs. S23**, mean firing rate, *n* = 13, two-sided Wilcoxon signed rank test, S-L: *P* = 0.97, L-S’: *P* = 0.92 and S-S’: *P* = 0.97; GS, S-L: *P* = 0.023, L-S’: *P* = 0.023 and S-S’: *P* = 0.86), rejecting the false positive discovery due to triangular node structures (**Figs. 3k-m** and **Supplementary Fig. S23**). The decrease of GS in large environments are consistent with the idea that grid cells anchor locally (Stensola et al., 2015). Overall, V2 grid cells exhibited similar features as those identified in the rat mEC (Hafting et al., 2005).

### Head-direction and conjunctive cells in V2

Head-direction (HD) cells discharge rapidly only when the animal’s head points towards a particular azimuth (direction) in the horizontal plane (Taube et al., 1990). We next investigated whether classical head-direction cells are present in the visual cortex. To categorize the preferred firing direction of V2 units, we used the mean vector length (MVL) to calculate the head directional tuning across 360°. A V2 unit was classified as a HD cell if its MVL exceeded the 99^th^ percentile of shuffled data (**Fig. 4b**). About 8.65% (92/1364) recorded V2 units met the significant criterion and had strong modulation to the animal’s heading direction (**Fig. 4a** and Supplementary Fig. 25). This percentage was significantly higher than expected by random selection from the entire shuffled population (**Fig. 4b**; Z = 28.40, *P* < 0.001; binomial test with expected *P*0 of 0.01 among large samples). We applied a maximum-likelihood correction procedure to alleviate the influence of spatial selectivity on biased HD sampling. Upon correction, about 77.97% (92/118) HD cells remained unchanged in their directional firing rate histograms (**Supplementary Fig. S24**), yet with 4.06% decrease in directional information; this was consistent with HD cells in the rat presubiculum and postsubiculum (Burgess et al., 2005). We further analyzed the remaining 92 V2 HD cells that passed the stringent 99^th^ percentile significance threshold, and these V2 cells had a mean firing rate of 2.09 ±0.22 Hz (**Fig. 4c**) and an average MVL of 0.58 ±0.06 (**Fig. 4d**). The preferred firing direction was stronger during running (> 2.5 cm/s) than immobility (< 2.5 cm/s) (**Fig. 4e** and Supplementary Fig. S9d, *n* = 92, *P* = 0.000, two-tailed paired *t*-test), yet the angular velocity between “preferred firing directions” and “non-preferred firing direction” was not significantly different (**Fig. 4f**, *n* = 92, *P* = 0.58, two-tailed paired *t*-test). The preferred firing direction of HD cell population uniformly distributed across 360° (**Fig. 4g**, *n* = 92, Rayleigh test for uniformity, *r* = 0.074, *P* = 0.61). The intra-trial directional selectivity remained stable between the first and second halves of individual recording sessions (**Fig. 4h** and Supplementary Figs. S10d-f). The full tuning width at half maximum (FWHM) of recorded V2 HD cells was 79.38° ± 8.27°, which was slightly greater than the statistic of presubicular HD cells but similar to the statistic in the lateral mammillary nuclei (**Fig. 4i** and **Supplementary Fig. S26**). Similar to those conjunctive cells reported in the rat mEC (Sargolini et al., 2006), we also found conjunctive place × HD cells (5/165 place cells), conjunctive border × HD cells (5/85 border cells) and conjunctive grid × HD cells (6/70 grid cells) in V2 (**Figs. 4j, k**).

**Fig 4.**
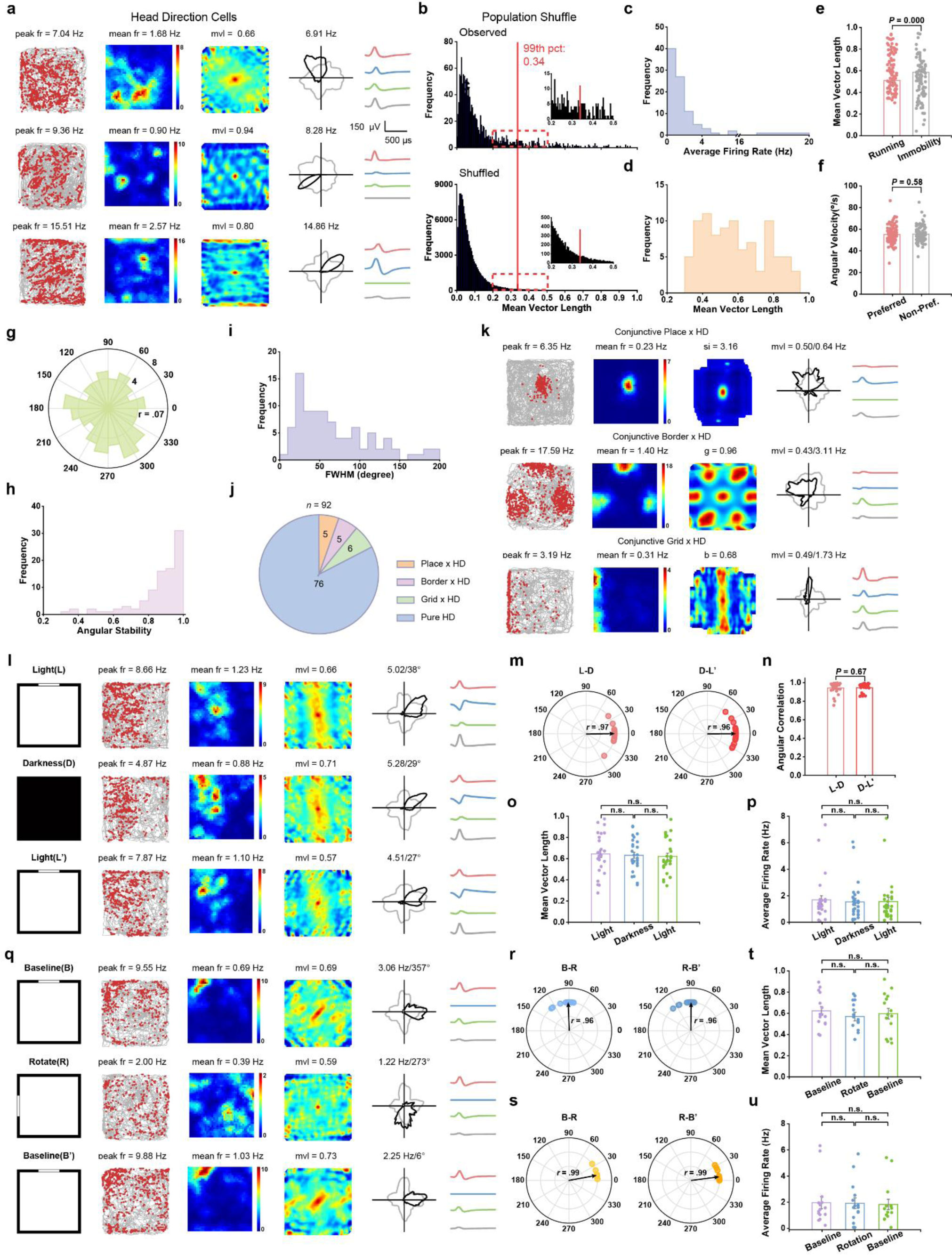
Head-direction (HD) and conjunctive cells in the visual cortex. a, Three representative V2 HD cells. Similar bol and notations as Fig. la. Mean vector length (MVL) for each HD cell is labelled at the top of firing rate map. Fourth column: HD tuning curves (black) plotted against dwell-time polar plot (grey). The directional plots show strong head directional tuning. **b**, Distribution of MVL in observed data (top panel) and shuffled data (bottom panel). Red line indicates the 99^th^ percentile for MVL derived from the shuffled data. The inset shows the zoomed-in of red dashed box. **c-d**, Distribution of mean firing rate and MVL for all V2 HD cells. **e**, HD tuning was stronger during running than slow movement or immobility (*n* = 92, two-tailed paired *t*-test, *P* = 0.000). **f**, Angular velocity was similar between the preferred and non-preferred firing directions for all V2 HD cells (*n* = 92, two-tailed paired *t*-test, *P* = 0.58). **g**, Polar plot showing the distribution of preferred direction for all HD cells (*n* = 92, Rayleigh test for uniformity: r = 0.074, *P* = 0.61). **h**, Distribution of intra-trial angular stability between the first and second halves derived from all HD cells (*n* = 92). **i**, Distribution of tuning widths defined as full width at half maximum (FWHM) of directional tuning curves derived from all HD cells (*n* = 92). **j**, Venn diagram showing the composition of conjunctive HD cells and pure HD cells. **k**, Representative conjunctive Place × HD, Border × HD and Grid × HD cells. **l**, Firing properties of one representative V2 HD cell in the light condition and darkness. **m**, Polar plot showing angular shifts in the preferred firing direction for all HD cells in the L-D-L’ conditions. V2 HD cells maintained their preferred firing directions in darkness (*n* = 28, L-D, Rayleigh test, *r* = 0.97, median shift = 3.49°; D-L’, Rayleigh test, *r* = 0.96, median shift = 7.44°; two-sided Wilcoxon signed rank test, *P* = 0.44 and 0.96 for D-L and L-D’, respectively). **n**, V2 HD cells showed high angular correlation in the L-D-L’ conditions (*n* = 28, two-sided Wilcoxon signed rank test, *P* = 0.67 between D-L and L-D’). **o**, MVL was not significantly different in the L-D-L’ conditions (*n* = 28, two-sided Wilcoxon signed rank test, *P* = 0.52, 0.47 and 0.36, respectively). **p**, Mean firing rates remained stable in the L-D-L’ conditions (*n* = 28, two-sided Wilcoxon signed rank test, *P* = 0.21, 0.67 and 0.41, respectively). **q**, Responses of one representative V2 HD cell under visual cue manipulation. Schematic of the experimental paradigms (left column); firing properties of V2 HD cells in baseline (upper panels), counterclockwi se 90°cue-rotation manipulation (middle panels) and back to baseline (lower panels). **r**, Firing properties of V2 HD cells following cue card rotation. Polar plot showing angular shifts in the preferred firing direction from light condition to total darkness (B-R, left) and from total darkness back to light condition (R-B’, right) for HD cells. V2 HD cells maintained their preferred firing directions in darkness (*n* = 15, B-R, Rayleigh test, *r* = 0.96, median shift = 92.23°; R-B’, Rayleigh test, *r* = 0.96, median shift = 89.63°; two-sided Wilcoxon signed rank test, *P* = 0.006 and 0.020 for B-R and R-B’, respectively). **s**, Same as panel **r** except for V2 HD cells that did not follow the cue card manipulation (*n* = 14, B-R, Rayleigh test, *r* = 0.96, median shift = 10.41°; R-B’, Rayleigh test, *r* = 0.96, median shift = 7.43°; two-sided Wilcoxon signed rank test, *P* = 0.48 and 0.06 for B-R and R-B’, respectively). **t**, MVL did not change by cue card manipulation (*n* = 15, two-sided Wilcoxon signed rank test, *P* = 0.10, 0.53 and 0.38, respectively). **u**, Mean firing rates were not significantly different between consecutive baseline, cue card rotation and baseline conditions (*n* = 15, two-sided Wilcoxon signed rank test, *P* = 0.78, 0.46 and 0.26, respectively).

We further tested the effect of visual landmarks on V2 HD cells during consecutive light-dark-light sessions (**Fig. 4l**). V2 head-direction cells maintained their preferred firing directions in the darkness (*n* = 28, Rayleigh test, for L-D: *r* = 0.97, median shift = 3.49°; for D-L’: *r* = 0.96, median shift = 7.44°; two-sided Wilcoxon signed rank test, *P* = 0.44 and 0.96 for D-L and L-D’, respectively, **Fig. 4m**). There was no significant change in angular correlation (*n* = 28, two-sided Wilcoxon signed rank test, *P* = 0.67 between D-L and L-D’, **Fig. 4n**), MVL (*P* = 0.52, 0.47 and 0.36, respectively, **Fig. 4o**) or mean firing rate (*P* = 0.21, 0.67 and 0.41, respectively, **Fig. 4p**), indicating that V2 HD tuning is not totally dependent upon visual inputs. About half (15/29) of V2 HD cells (**Fig. 4q** and **Supplementary Fig. S27**) produced a nearly equal shift in preferred firing direction (*n* = 15, Rayleigh test: for B-R, *r* = 0.96, median shift = 92.23°; for R-B’, *r* = 0.96, median shift = 89.63°; two-sided Wilcoxon signed rank test, *P* = 0.006 and 0.002 for B-R and R-B’, respectively, **Fig. 4r**). V2 HD cells also anchored to salient visual cues with no significant change in MVL (*n* = 15, two-sided Wilcoxon signed rank test, *P* = 0.10, 0.53 and 0.38, respectively, **Fig. 4t**) or mean firing rate (*P* = 0.78, 0.46 and 0.26, respectively, **Fig. 4u**). However, about half (14/29) of V2 HD cells did not follow the rotated cue card (*n* = 14, Rayleigh test: for B-R, *r* = 0.96, median shift = 10.41°; for R-B’, *r* = 0.96, median shift = 7.43°; two-sided Wilcoxon signed rank test, *P* = 0.48 and 0.06 for B-R and R-B’, respectively, **Fig. 4s**). Put together, V2 HD cells shared similar characteristics of other HD cells in the hippocampal-entorhinal network and the presubiculum (Taube, 2007).

## Discussion

The brain is responsible for transforming and integrating multimodal sensory information in the environment into neural representations that can be used for perception and action. The classical hippocampal-entorhinal network is fundamental to guide spatial navigation, displaying distinct spatially tuned cells in rodents, bats and nonhuman primates (Buzsaki and Moser, 2013). Sensory input and movement are coupled in real world, and it remains unclear how sensory input provides a complementary spatial or cognitive map while navigating in a naturalistic environment. Recently, strong spatial tuned neurons have been found in the retrospelinal cortex (RSC) (Alexander and Nitz, 2017; van Wijngaarden et al., 2020), posterior parietal cortex (PPC) (Whitlock et al., 2012; Wilber et al., 2014), and primary somatosensory cortex (Long and Zhang, 2021). Vision supports spatial navigation by providing distal visual cues and identifying visual landmarks (Flossmann and Rochefort, 2020; Nau et al., 2018; Saleem, 2020). It has been shown that visual thalamic and cortical neurons are modulated by spatial signals in both virtual and naturalistic environments (Diamanti et al., 2021; Fiser et al., 2016; Fournier et al., 2020; Hok et al., 2018; Jankowski and O’Mara, 2015; Leinweber et al., 2017; Pakan et al., 2018; Saleem et al., 2013; Saleem et al., 2018; Taube, 1995; Vantomme et al., 2020). Spatial representations discovered in multiple visual areas is not surprising since the visual system is responsible for transforming sensory information from eye-centered to world-centered coordinates and estimating self-location (Fournier et al., 2020). Visual cues may provide a bias for position estimate based on non-visual cues (such as distance or speed) in the context of predictive coding(Acharya et al., 2016; Rao and Ballard, 1999; Stachenfeld et al., 2017). Although many studies focused on how visual information influences spatial tunings of the hippocampal-entorhinal network or whether responses of visual neurons are transformed from the hippocampal-entorhinal network, the neural basis of the spatial navigation system in visual cortex remains unclear. Our study provides the first identification and systematic characterization of four functionally distinct spatial cell types in V2, which may form the basis of a compact visual spatial navigation system. These diverse spatial tunings resemble those reported in the classical hippocampal-entorhinal network (Buzsaki and Moser, 2013) and identified recently in the rat primary somatosensory cortex (Long and Zhang, 2021).These recent and current findings challenge the traditional view, and suggests the cognitive maps are more widely represented in the brain than traditionally thought (O’Mara and Aggleton, 2019); **Supplementary Fig. S28**).

The dLGN, being the one synapse upstream structure of the V1, displays place cells (Hok et al., 2018). Multiple lines of evidence have shown that V1 neurons are also modulated by animal’s spatial position (Diamanti et al., 2021; Fiser et al., 2016; Flossmann and Rochefort, 2020; Fournier et al., 2020; Haggerty and Ji, 2015; Ji and Wilson, 2007). Calcium imaging in head-fixed mice in a virtual reality environment demonstrated that spatial modulation in V1 strengthens with experience as well as with active behavior (Diamanti et al., 2021). The V2, being the downstream structure of V1, have been studied in spatial navigation tasks with reported 5% egocentric boundary cells (Gofman et al., 2019). However, their implanted electrode location was closer to the postrhinal cortex (POR) than V2, which may explain the difference between their findings and ours. Additionally, the medial and lateral PPC display spatial coding (Nitz, 2006; Save and Poucet, 2009; Whitlock et al., 2012; Wilber et al., 2014). As the rat PPC is located rostral to the primary and secondary visual cortical areas, and caudal to the somatosensory cortex (∼3.5-5 mm posterior of bregma), the exact location of the implanted electrodes may explain the difference in experimental findings (Gofman et al., 2019; Whitlock et al., 2012).

An important question remains: where do the V2 spatial and directional signals come from? In the visual hierarchy, the V2 receives strong feedforward connections from V1, and sends strong connections to higher visual cortical areas as well as feedback connections to V1. The role of V2 in visual cortex has been widely investigated, splitting the dorsal and ventral streams. The ventral visual-to-hippocampal stream is known to play an important role in visual memory or object-recognition memory (Bussey and Saksida, 2007; Lopez-Aranda et al., 2009). Feedback signals from the classical hippocampal-entorhinal GPS network might contribute to the source of non-visual navigational signals in V2. There are reciprocal projections between V2 and several parahippocampal areas including the perirhinal cortex (PER), POR, RSC, MEC, lateral entorhinal cortex (LEC), presubiculum (PrS) and parasubiculum (PaS) (Olsen et al., 2017), which have been reported to encode both egocentric and allocentric spatial and directional information (Alexander et al., 2020). Although there is a lack of direct intrinsic projections between V2 and hippocampus, spatial and directional representations of V2 might be influenced by the hippocampal spatial information through indirect cortico-cortical connections such as the RSC, which is known to encode cortical spatial and directional signals (Mao et al., 2017; Mao et al., 2018; van Strien et al., 2009). Development of large-scale multi-site *in vivo* electrophysiological recording techniques may prove crucial to provide a complete picture of sensory coding of visual systems (Buzsaki et al., 2015; Koay et al., 2020; Siegle et al., 2021). Converging evidence has shown that higher-order visual areas such as V2 may encode additional non-visual cognitive cues including the location, direction and motion. This is also consistent with spatial modulation signals being observed only in higher visual cortices but not in lower visual cortices along the visual hierarchy (Diamanti et al., 2021). We envision that the visual cortex encodes a compact spatial map in parallel to the hippocampal-entorhinal network, and provides a complementary multimodal cognitive mapping of the external world. Just like the hippocampal neurons can jointly encode space and time (Buzsaki and Moser, 2013; Eichenbaum, 2014), many sensory cortical neurons may have mixed selectivity in complex tasks (Siegle et al., 2021; Steinmetz et al., 2019). It is very likely that the spatial modulation is an omnipresent property of pyramidal neurons across many sensory cortices (Hawkins et al., 2018).

It remains to be determined whether these high-order visual areas form the unique neural basis of spatio-visual map. A causal circuit dissection based on optogenetic inactivation of individual visual areas along the visual thalamocortical pathway may help answer this question in the future (Buzsaki et al., 2015; Zhang et al., 2013). It is important to distinguish the visuospatial or spatio-visual map in the visual system between active sensing (e.g., active vision and active navigation) and passive sensing (e.g., purely visual exploration), or between naturalistic and virtual reality environments. Proprioceptive feedback and sensorimotor integration may play a vital role in reshaping spatial selectivity of hippocampal and cortical neurons (Acharya et al., 2016; Aghajan et al., 2015; Ravassard et al., 2013). It may not come as a surprise that the spatio-visual map in visual cortex encoded both egocentric and allocentric spatial cues in parallel (Bicanski and Burgess, 2020; LaChance et al., 2019; Wang et al., 2020). How and whether hippocampal-entorhinal spatial signals interact with non-visual navigational signals in the visual cortex awaits for further investigation.

## Materials and Methods

### Long-Evans Rats

The recording experiments were performed on six male Long-Evans adult rats (2-4 months) weighting 250-450 grams on the day for chronic surgery. Rats were individually housed in transparent plastic cages (W × L × H: 35 cm × 45 cm × 45 cm) and kept on a 12-hour reversed light/dark schedule (lights on from 21:00 p.m. to 09:00 hours). All behavioral trials were performed during the dark phase. Rats were maintained in a temperature-controlled (19-23°C) and humidity-adjusted (55-70%) vivarium.

Rats were partially food-deprived to about 85-90% of free-feeding body weight. Food restriction was imposed 8-24 hours before each training and recording trial. Water was supplied *ad libitum*. All animal experiments were performed in accordance with the National Animal Welfare Act under a protocol with the permission license number #SYXK-2017002 approved by the Animal Care and Use Committee from both the Army Medical University and Xinqiao Hospital.

### Surgical procedures and tetrode placement

The rats were anesthetized with isoflurane mixed with oxygen (1.5-3.0% in O2), immobilized in a stereotaxic frame (David Kopf Instruments, Tujunga, California, USA) and kept on feedback-adjusted temperature control pad at 37°C. A self-assembled microdrive loaded with four tetrodes were implanted to target the mediolateral (V2ML) and mediomedial (V2MM) regions of the second primary cortex with the stereotaxic coordinate centered at ∼2.6 mm lateral to the midline (ML), ∼−4.7 mm anterior-posterior (AP) from bregma, 0.4-1.1 mm dorsal-ventral (DV) below the dura and at an angle of 10°from the medial-to-lateral direction in the coronal plane, secured with dental cement with 8-10 anchor screws. A screw served as the ground electrode. Tetrodes were assembled with four 17 µm polyimide-coated platinum-iridium (90-10%) wires (#100167, California Fine Wire Company). Tetrodes were plated with a 1.5% platinum solution to reduce their impedances between 150 and 300 kΩ at 1 kHz through electroplating prior to surgery (nanoZ; White Matter LLC, Seattle, Washington, USA). 8-10 jeweler screws were threaded into the rat skull, and individual microdrives were anchored to screws with several rounds of application of the dental cement.

### Behavioral protocol and data collection

The chronically implanted rats were allowed at least one week of recovery before behavioral training, tetrode turning and data recording were initiated. Rats were trained to forage around in a 1 × 1 m^2^ square box with a white cue card (297 × 210 mm^2^) mounted on one side of the interior wall. Food pellets were randomly dispersed into the enclosure intermittently to encourage free exploration.

Each recording trial lasted typically between 20 and 40 min to facilitate full coverage of the testing enclosure. Tetrodes were lowered very slowly in steps of 25 or 50 µm daily until well-separated single units can be identified. Data were acquired by an Axona system (Axona Ltd., St. Albans, U.K.) at 48 kHz, band-passed between 0.8-6.7 kHz and a gain of x5-18k. Spikes were digitized with 50 8-bit sample windows. Local field potentials were recorded from one of the electrodes with a low-pass filter (cut-off: 500 Hz).

### Spike sorting, cell-type identification and locational firing rate map

To identify well-isolated units, spike sorting was manually performed offline with graphical cluster-cutting software (TINT, version 4.4.16, Axona Ltd, St. Albans, U.K.), and the clustering was primarily based on features of the spike waveform (peak-to-trough amplitude and spike width), together with additional autocorrelations and cross-correlations separation tools. During the manual cluster cutting, we strictly counted units with similar or identical waveform shapes only once whenever similar or identical individual units were recorded and tracked across two consecutive recording sessions. Only recording sessions in which rats covered more than 80% of the running environment were taken for further analysis. To assess the quality of cluster separation, we calculated the well-established metric (Lratio) that measures the isolation distance between clusters (Schmitzer-Torbert et al., 2005).

Two small light-emitting diodes (LEDs) were mounted to the headstage to track the rats’ speed, position and orientation via an overhead video camera. Only spikes recorded during animal’s instantaneous running speeds > 2.5 cm/s were chosen for further analysis in order to exclude confounding behaviors such as immobility, grooming and rearing.

To classify firing fields and firing rate distributions, the position data were divided into 2.5 × 2.5 cm^2^ bins, and the path was smoothed with a 21-sample boxcar window filter (400 ms; 10 samples on each side). Units with > 100 spikes per session and with a coverage of >80% were included for further analyses. Maps for spike numbers and spike times were smoothed with a quasi-Gaussian kernel over the neighboring 5 × 5 bins. Spatial firing rates were calculated by dividing the smoothed map of spike numbers with spike times. The peak firing rate was defined as the highest rate in the corresponding bin within the spatial firing rate map. Mean firing rates were averaged from the whole session data. The criterion for classifying a V2 unit as a spatially tuned cell (e.g., place cell, head-direction cell, border cell or grid cell) was determined by measuring whether the corresponding score was greater than the 99^th^ percentile value of the shuffled data.

### Identification of place cells

Spatial information (SI) is a quantification of the extent to which a unit’s firing pattern can predict the position of freely moving animals and is expressed in the unit of bits per spike:

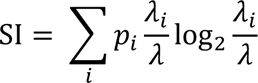

where λ_*i*_ is the mean firing rate of the unit in the i-th bin, λ is the overall mean firing rate of the unit in the trial, and *p*_*i*_ is the probability for the animal being at the location of the i-th bin. Adaptive smoothing was applied to optimize the trade-off between spatial resolution and sampling error before the SI calculation. The data were first divided into 2.5 × 2.5-cm^2^ bins, and then the firing rate within each bin was calculated by expanding a circle centered on the bin until

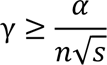

where γ is the circle’s radius in bins, n is the number of occupancy of samples within the circle, s is the total number of spikes fired within the circle, and α =10000 is a constant. With a position sampling frequency at 50 Hz, the firing rate assigned to that bin was then set to 50 · *s*/*n*.

A V2 unit was classified as a place cell if its SI was above the chance level, which was computed by a random permutation process using all recorded cells. For each round of the shuffling process, the entire sequence of spike trains from each unit was time-shifted along the animal’s trajectory by a random period between 20 s after the trial onset and the trial duration minus 20 s, with the end wrapped to the beginning of the trial. A spatial firing rate map was then constructed, and spatial information was calculated. This shuffling process was repeated 100 times for each unit, generating a total of 136,400 permutations for the 1364 V2 units. This shuffling procedure was aimed to preserve the temporal firing characteristics in the unshuffled data while disrupting the spatial structure at the same time. The SI score was then measured for each shuffled rate map. The distribution of SI scores across all 100 permutations of all units was computed, from which the 99^th^ percentile of the significant level was determined. To determine the threshold for categorizing a V2 unit as a place cell, we chose the SI score above the 99^th^ percentile of the distribution from shuffled populations. In addition to population shuffling, within-unit shuffling was performed within the entire sequence of spike trains from each individual unit, and the shuffling process was repeated 100 times for each single unit.

Spatial sparsity was used to measure how compact and selective the place field is relative to the recording enclosure. The spatial sparsity was calculated using the formula as follows:

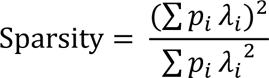

where *p*_*i*_ is the occupancy probability for the animal being at the location of the i-th bin in the map and λ_*i*_ is the mean firing rate of the cell in bin i.

Spatial coherence was estimated by calculating the mean correlation between the firing rate of each bin in the map and the aggregate firing rate of the eight nearest bins. We used the unsmoothed firing rate maps for computing the spatial coherence.

The spatial correlation across trials from the same recording arena was computed for each cell by correlating the firing rates in corresponding paired bins of two smoothed rate maps. The spatial stability within trials was estimated by calculating spatial correlations between firing rate maps generated from the first and second halves of the same trial. Place cells with spatial stability lower than 0.3 were excluded for further analysis.

### Identification of grid cells

Gridness score (GS) was previously established to quantify the grid cell (Boccara et al., 2010; Hafting et al., 2005; Zhang et al., 2013). We calculated the spatial autocorrelation with smoothed rate maps. The unbiased autocorrelograms were derived from Pearson’s product-moment correlation coefficient correcting for edge effects and unvisited locations.

Let λ(*x,y*) denote the unit’s mean firing rate at a two-dimensional Cartesian coordinate (*x,y*), the autocorrelation of the spatial firing field was calculated as:

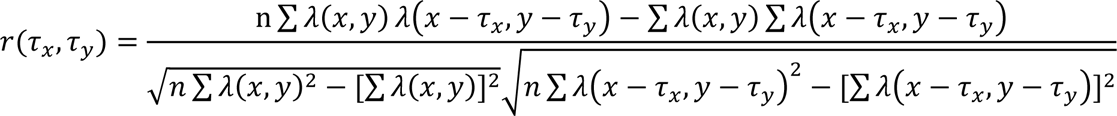

where the summation was over n pixels for both λ(*x*,*y*) and λ(*x*−τ_x_, *y*−τ_y_) (τ_x_ and τ_y_ denote the spatial lags). Autocorrelations were not calculated for spatial lags of τ*_x_*, τ_y_ where n < 20.

The degree of spatial regularity (“gridness” or GS) was calculated for each unit by using a circular sample centered on the central peak of the autocorrelogram but excluding the central peak itself, and by comparing rotated versions of this circular sample. Pearson’s correlations between this circular sample and its rotated versions were calculated, with the angles of rotation of 60°and 120°in the first group, and 30°, 90°and 150°in the second group. A unit’s GS was defined as the minimal difference between any of the coefficients in the first group and any of the coefficients in the second group. Shuffling was performed in the same procedure used for defining place cells. Grid cells were defined if their rotational symmetry-based GSs exceeded the 99^th^ percentile of the distribution of GS derived from the shuffled data.

Grid spacing was computed as the median distance from the grid center to the closest peak among six neighboring firing fields in the autocorrelogram of spatial firing map. Since this analysis might be sensitive to noise in the grid autocorrelogram, grid spacing was computed only when the median distance to the six neighboring peaks for the analyzed unit was comparable to the radius of the circle centered on the gird autocorrelogram with the highest GS. The radius of this circle around the center of autocorrelogram was also referred to as the grid field size.

Grid orientation was calculated by first computing vectors from the center of the autocorrelogram to each of the three adjacent peaks among six neighboring firing fields in the autocorrelogram of spatial firing map in the counterclockwise direction, beginning with a camera -based reference line of zero degree. The angle between the minimal orientation of those three vectors and the camera - based reference line was defined as the grid orientation.

### Identification of head-direction cells

The calculation of mean vector length (MVL) was followed by previous publication (Boccara et al., 2010; Sargolini et al., 2006; Zhang et al., 2013). The rat’s head-direction was computed by the relative position of two LEDs differentiated through their sizes. The directional tuning curve for each recorded unit was drawn by plotting the firing rate as a function of the rat’s head angle, which is divided into bins of 3° and then smoothed with a 14.5° mean window filter (2 bins on each side). To avoid potential biases, data were only used if all head angle bins contained nonzero values.

The strength of directionality was measured by computing the MVL from circular distributed firing rates. The chance value was determined by a shuffling process simulated in the same way as for place cells, with the entire sequence of spike trains time-shifted between 20 s and the whole trail length minus 20 s along the animal’s trajectory. A V2 unit was defined as a head direction cell if its mean vector length was larger than the 99^th^ percentile of MVLs in the shuffled distribution. Angular stability was computed by calculating the correlation of firing rates across directional bins generated from the first and second halves of the same trial. Head direction cells with angular stability lower than 0.3 were excluded for further analysis.

### Identification of border cells

The calculation of the border score (BS) was followed by previously literature (Boccara et al., 2010; Solstad et al., 2008; Zhang et al., 2013). Border or boundary vector cells were identified by calculating the border score, or the difference between the maximal length of any of the four walls touching on any single spatial firing field of the cell and the average distance of the firing field to the nearest wall, divided by the sum of those two values. The BS ranged from −1 for cells with perfect central firing fields to +1 for cells with firing fields that exactly line up with at least one entire wall. Firing fields were defined as summation of neighboring pixels with total firing rates higher than 0.3 times the unit’s maximum firing rate that covered a total area of at least 200 cm^2^.

Border cell classification was verified in the same way as for place cells, grid cells, and head direction cells. For each permutation trial, the whole sequence of spike trains was time-shifted along the animal’s trajectory by a random period between 20 s and 20 s less than the length of the entire trial, with the end wrapped to the start of the trial. A spatial firing rate map was then obtained, followed by BS estimation.

The distribution of BS was calculated for the entire set of permutation trials from all recorded units, from which the threshold with the 99^th^ percentile was determined. A V2 unit was defined as a border cell if its BS was higher than the 99^th^ percentile threshold derived from the shuffled data.

### Correction procedure for inhomogeneous sampling

To assess the inhomogeneous sampling bias on the spatial firing properties of V2 place cells and head-direction cells, we applied a maximum-likelihood correction procedure (originally proposed by Burgess and colleagues, and codes generously shared by the authors) (Burgess et al., 2005). Briefly, the position data were sorted into 2.5 × 2.5 cm^2^ bins and directional date into 120 bins of 3°each before applying the correction algorithm. The field plot was applied with 5 boxcar × 5 boxcar smoothing, and polar plot was applied with 5 boxcar smoothing for the purpose of visualization.

### Environmental manipulations

For visual landmark rotation, we first recorded V2 unit activity in the standard recording session followed by a 90°cue-card rotation in the clockwise or counterclockwise orientation. Next, another standard session was performed with the cue-card rotated back to the original position. For recording in the elevated platform without walls, we first recorded V2 units in the square box, followed by the recording in the elevated platform without walls. Finally, the animal returned to the original square box for another recording session. For the recording of border cells in the presence of inserted wall, we first identified V2 border cells in the square box. Then the recording session was followed by the insertion of a wall along the center of the external wall. Another recording session was performed after removing the inserted wall. For food/no food comparisons, we recorded V2 units in two consecutive recording sessions without and with throwing food pellets into the running enclosure.

### Histology and electrode track location

After completion of the final recording sessions, implanted rats were deeply anaesthetized with sodium pentobarbital (0.01ml/g) and perfused transcardially with ice-cold 1 x phosphate-buffered saline (PBS) followed by 4% ice-cold paraformaldehyde (PFA) in 1 x PBS. Brains were removed and post-fixed in 4% PFA in 1 x PBS at 4°C overnight. The brain was then transferred into 10, 20 and 30% sucrose/PFA solution sequentially across 72 hours before sectioning by using a cyrostat. Thirty-micron-thick coronal sections were serially cut and obtained through the implanted brain area. Sections were mounted on glass slides and stained with Cresyl Violet (Sigma-Aldrich). The final recording positions were imaged and determined from digitized images of the Nissl-stained sections scanned on the Olympus Slideview VS200 Digital Slide Scanner. Positions of each individual recordings were estimated from the deepest tetrode track according to the daily notebook on tetrode advancement. The tissue shrinkage correction was calculated by dividing the distance between the brain surface and electrode tips by the final advanced depth of the recording electrodes. Electrode traces were confirmed to be located between the mediolateral part of the secondary visual cortex (V2ML) and mediomedial part of the secondary visual cortex (V2MM) from five implanted rats based on the reference figures (from Figure 68 to 73) published in the sixth edition of *The Rat Brain in Stereotaxic Coordinates* (Paxinos and Watson, 2007).

## Data availability

Recording dataset will become available in a forthcoming public domain, and reasonable request into acquiring the raw data beforehand should be directed to the corresponding author.

## Code availability

The source codes used in this study are available from the corresponding author upon reasonable request.

## Acknowledgments

We thank Hao Wu, Wan-Neng Tang, Jia-Shun Ren, Cai-Ping Song, De-Guang Qi, Tong-Quan Liao, Hao Chen, Qian Chen, Yang Yang, Hui Yang and Sheng-Qing Lv for their encouragement and support. We thank Edvard Moser, May-Britt Moser, Neil Burgess and Caswell Barry for sharing their analytical codes with us. Special thanks to Ting-Ting Huang, Mi Zhang and A- Xiang Zhou for their technical assistance. S.-J.Z. was supported by the National Natural Science Foundation of China (Grant# 31872775). X.L. is supported by the Chongqing Municipality postdoctoral fellowship (Grant# cstc2019jcyj-bshX0035). Z.S.C. is partly supported by the US National Institutes of Health (R01-MH118928).

## Author contributions

S.-J.Z. conceived the project. X.L. and S.-J.Z. designed the study, performed the surgery and analyzed the data. J.C., B.D. and X.L. conducted the experiments and acquired the recordings. Z.S.C. and B.D. provided the feedback. X.L. developed the codes and made the figures. Z.S.C., X.L. and S.-J.Z. wrote the manuscript.

## Additional information

### Supplementary information

(Supplementary **Figs. S1**-**S28**) accompanies this paper.

## Supplementary Materials

### Supplementary Fig. S1-S28

**Fig. S1.**
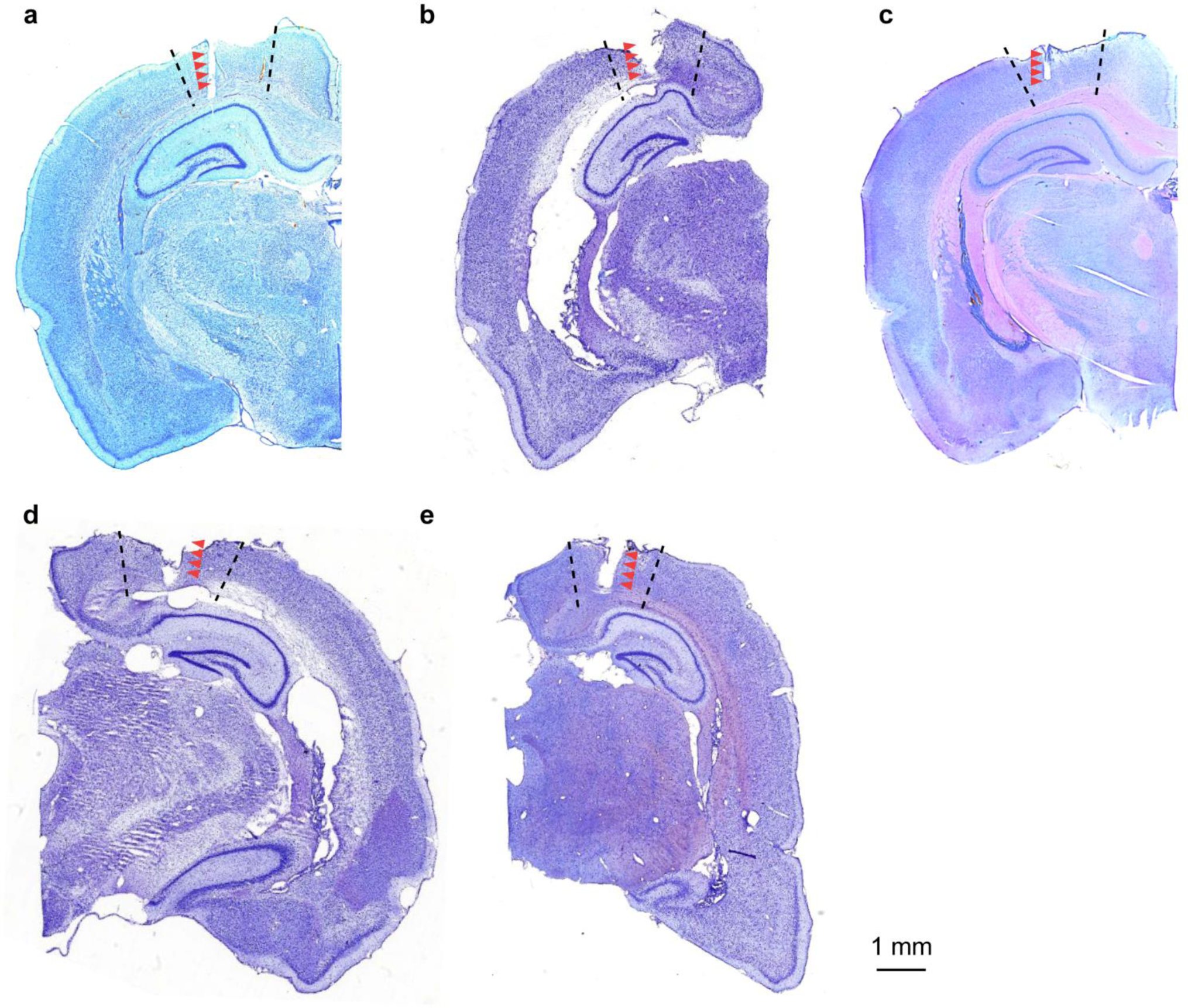
Tetrode recording sites in the rat secondary visual cortex (V2). **a-e**, Cresyl violet-stained coronal brain sections showing the electrode track and recording locations (red arrows) in V2 from five rats. Dashed lines indicate the boundaries of V2.

**Fig. S2.**
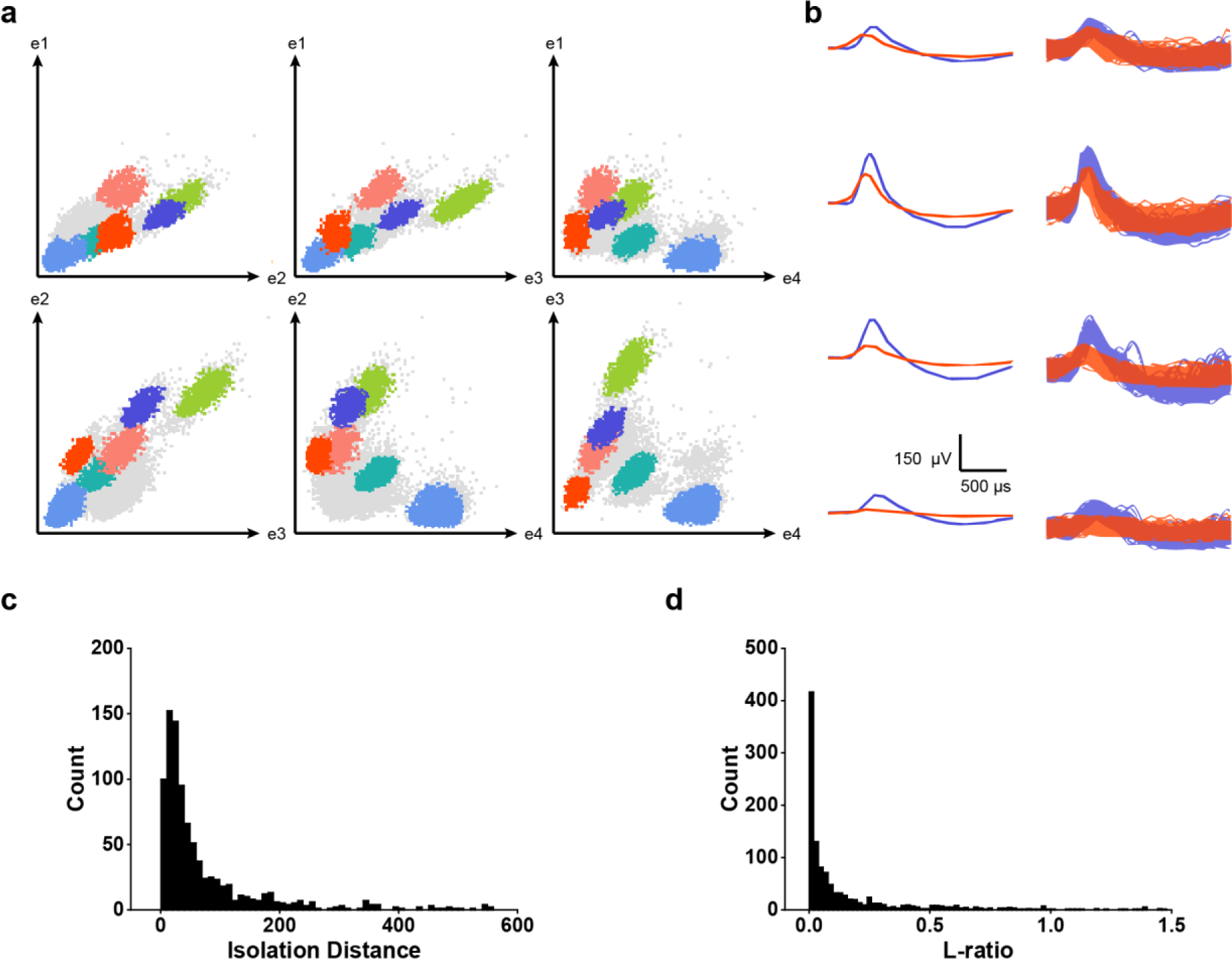
Cluster diagrams and isolation quality of spikes recorded from V2. **a**, Scatterplots showing the distribution of the peak-to-trough amplitudes for all spikes recorded in four electrodes of the same tetrode. **b**, Overlaid waveforms from two well-separated red and cyan-blue clusters in the corresponding scatterplot. **c-d**, Isolation Distance and L-ratio for recorded V2 units, respectively.

**Fig. S3.**
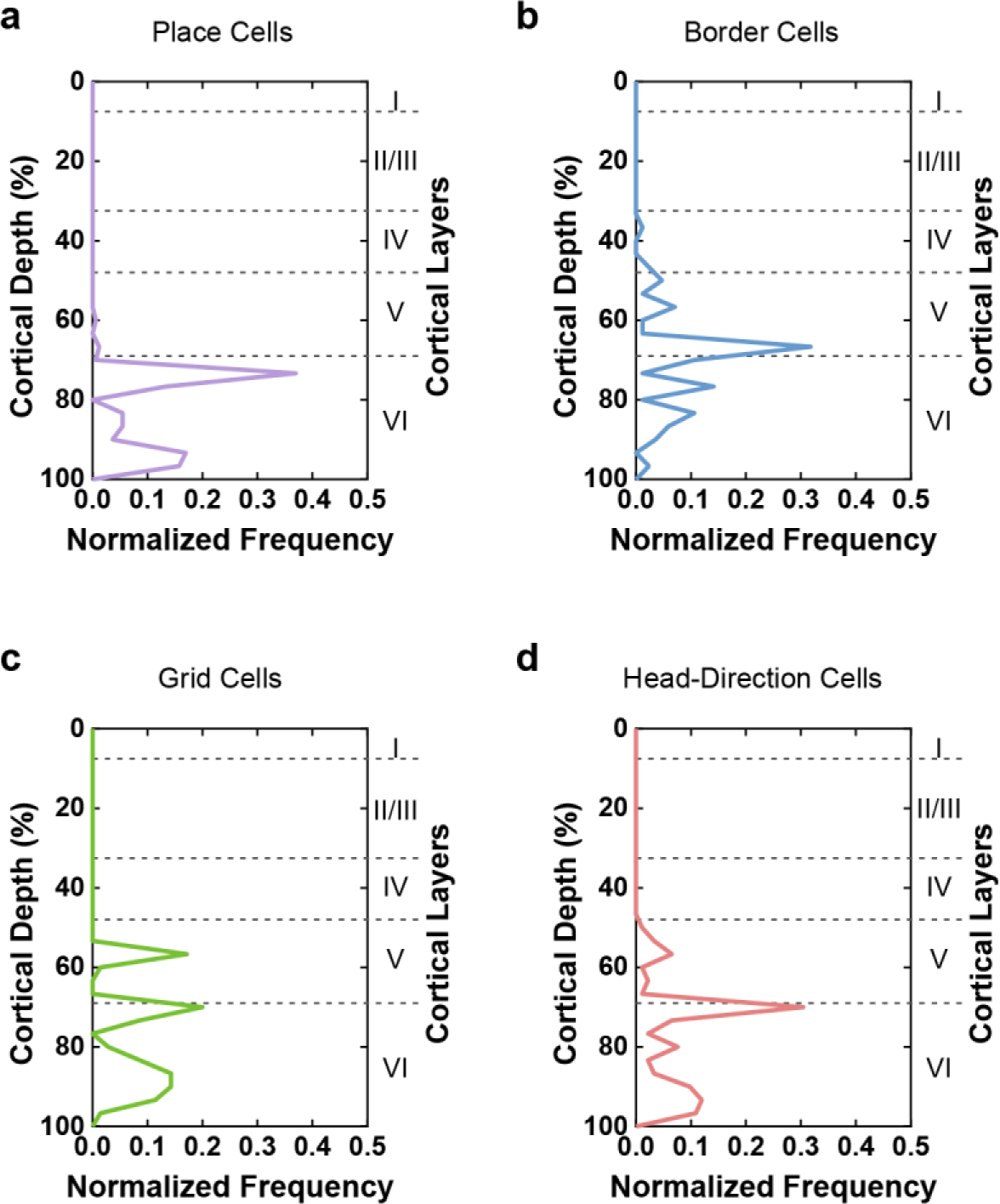
Layer distribution of four distinct V2 spatially tuned cell types. **a-d**, Normalized layer distributions of place cells, border cells, grid cells and head-direction cells in the rat V2, respectively. Note the near absence of recording in the superficial layer I-III.

**Fig. S4.**
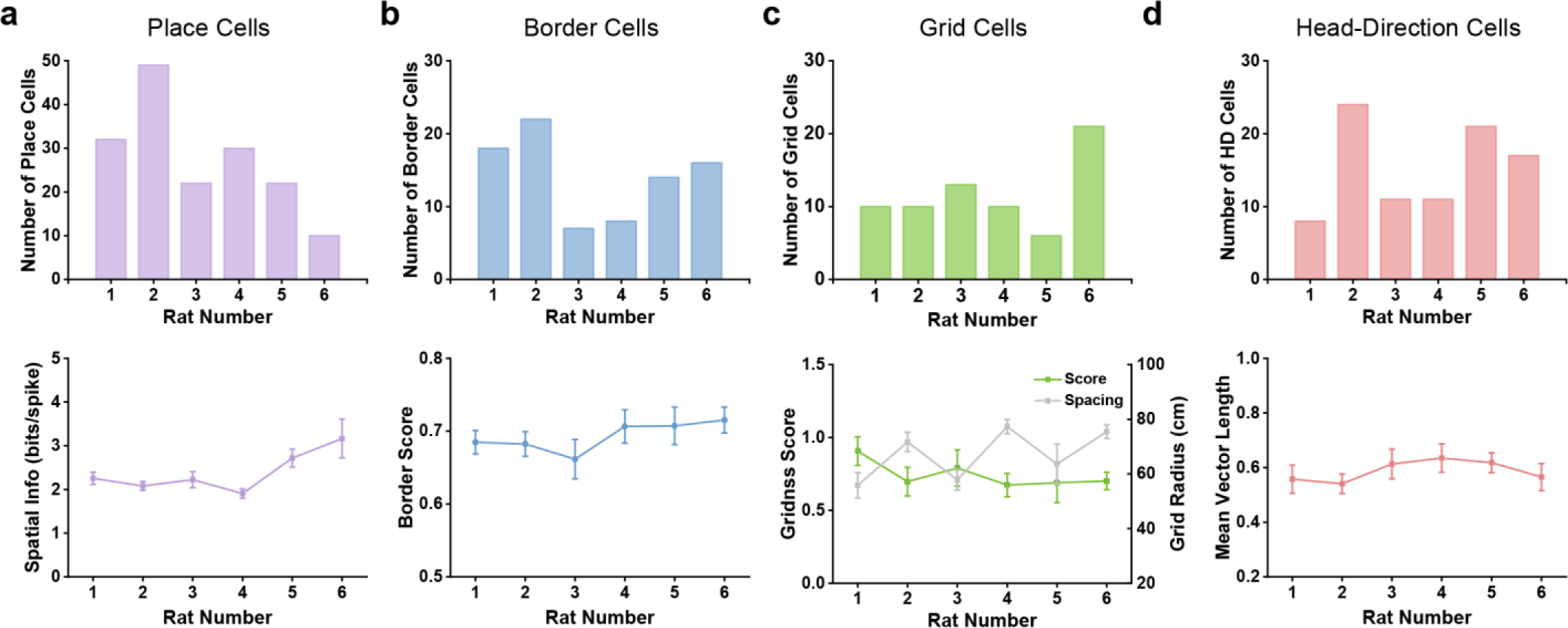
Spatial characterizations of recorded spatially tuned V2 units across six rats. **a-d**, Number of four different spatial cell types recorded in each rat (upper panels) and five spatial properties (spatial information, border score, gridness score, grid radius, and mean vector length) across different rats (bottom panels).

**Fig. S5.**
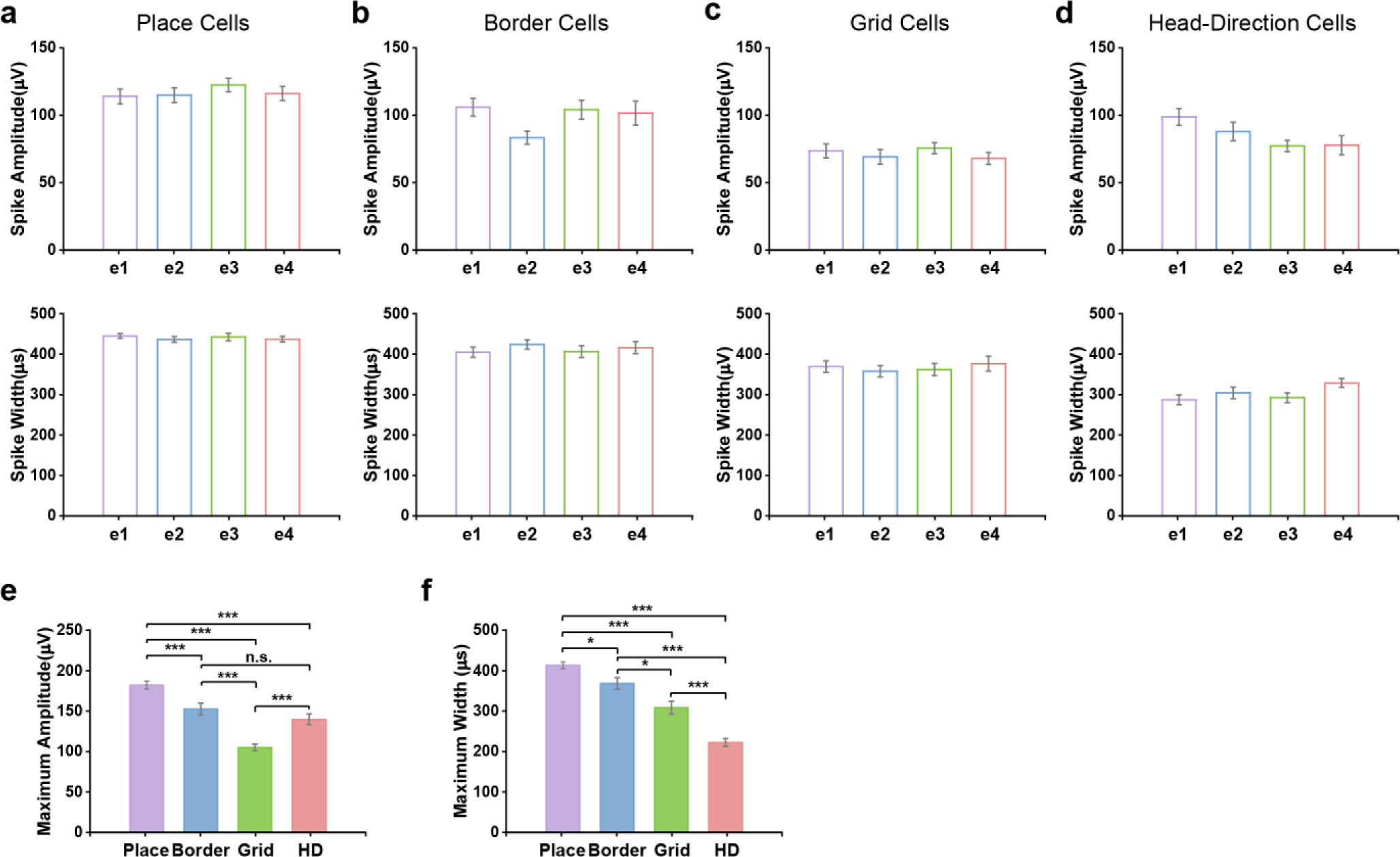
Distribution of spike amplitudes and spike widths for four types of V2 spatially tuned units. **a-d**, Peak-to-peak amplitudes (upper panels) and peak-to-trough widths (bottom panels) of spike waveforms on four electrodes (e1-e4) of the same tetrode for identified V2 place cells, border cells, grid cells and head-direction cells, respectively. **e-f,** Comparison of the highest amplitude on four electrodes and corresponding spike width for four distinct V2 spatially tuned cell types (*n* = 165, 85, 70 and 92, respectively, unpaired *t*-test, n.s., *P*>0.05; *, *P*<0.05; **, *P*<0.01; ***, *P*<0.001).

**Fig. S6.**
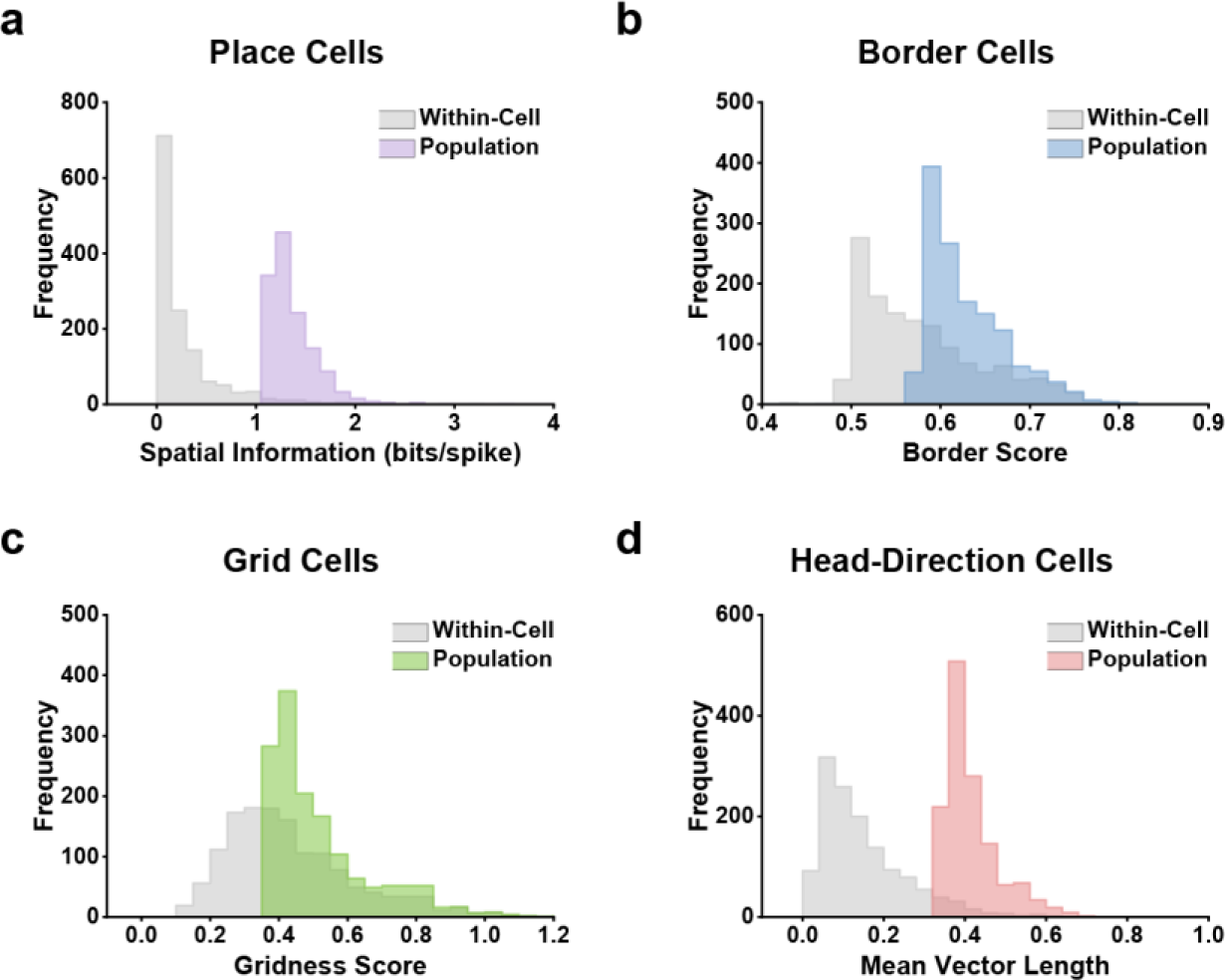
Comparison of threshold statistics determined by population shuffling and within-cell shuffling. **a-d**, Histograms showing the 99^th^ significance level of each randomly shuffled distribution by population shuffling and within-cell shuffling to calculate the spatial information, border score, gridness score and mean vector length for all identified V2 units, respectively.

**Fig. S7.**
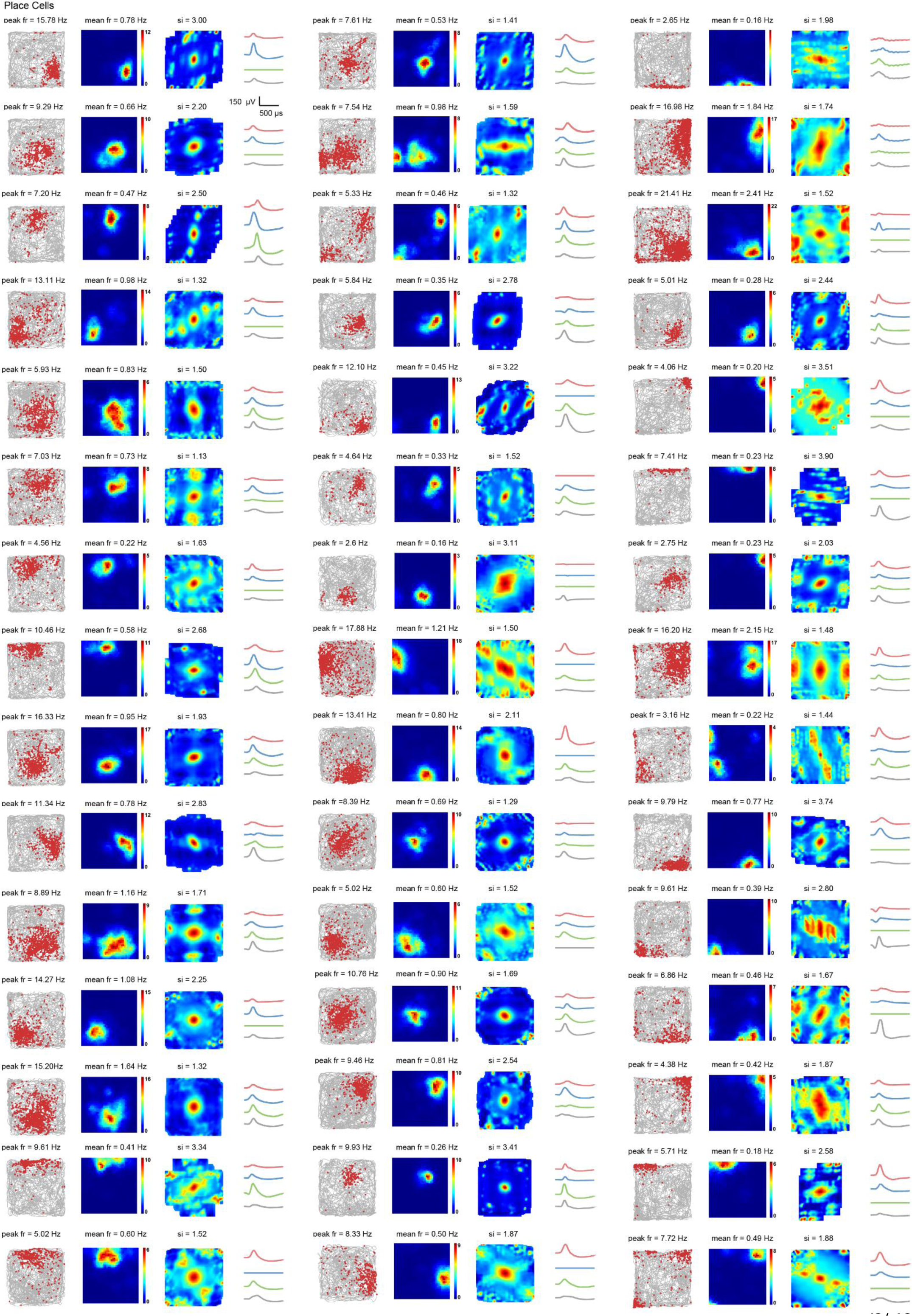
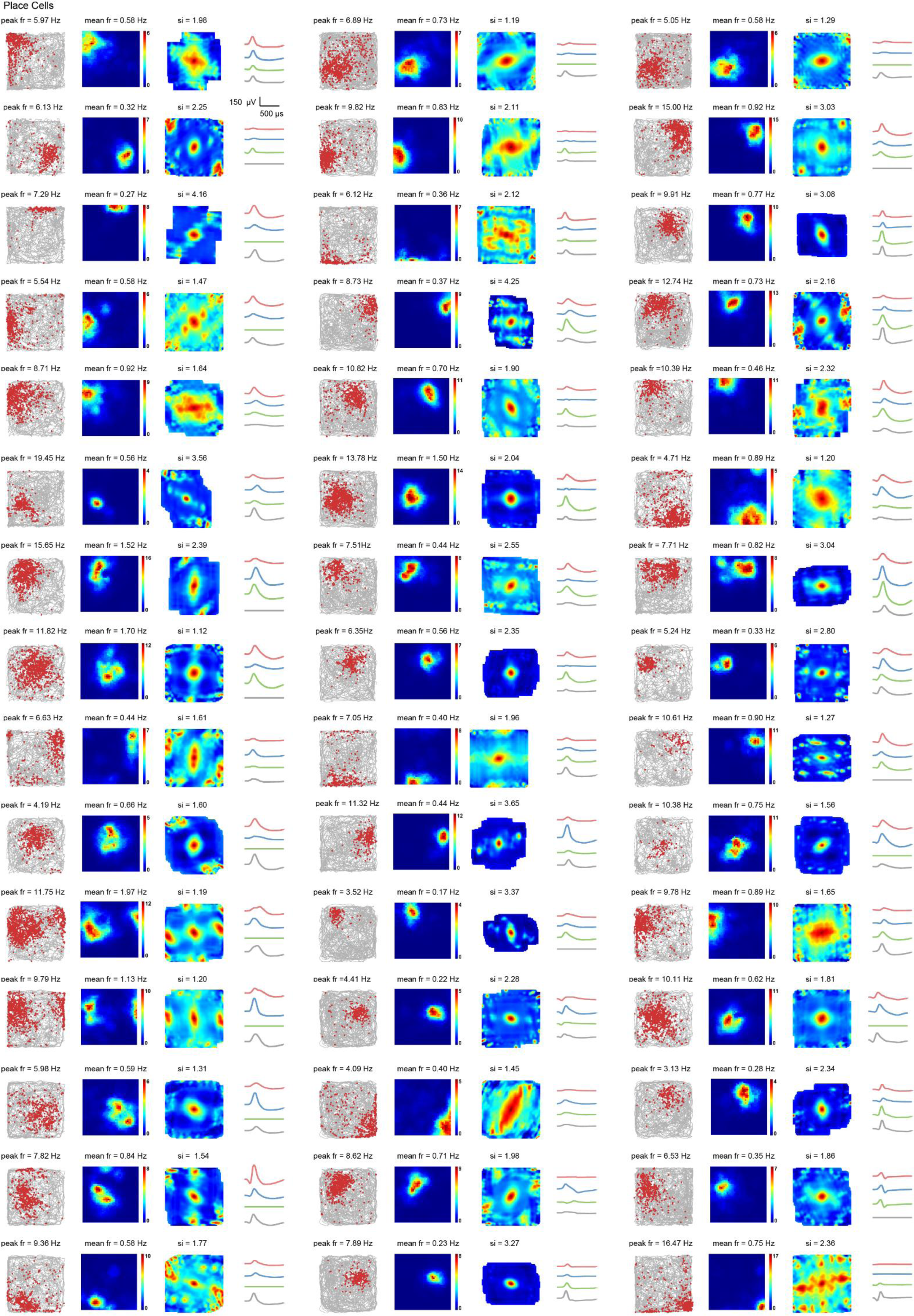
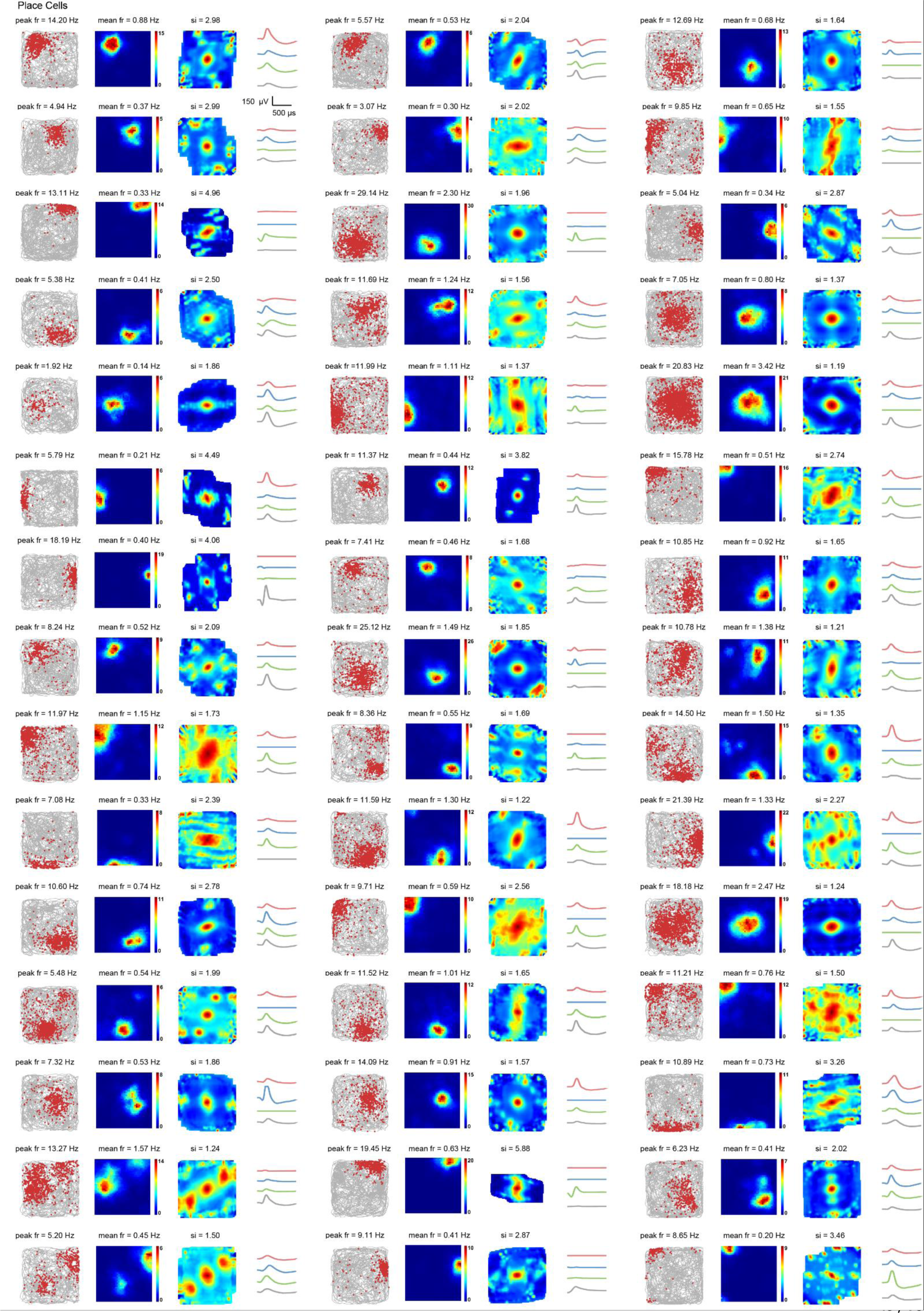
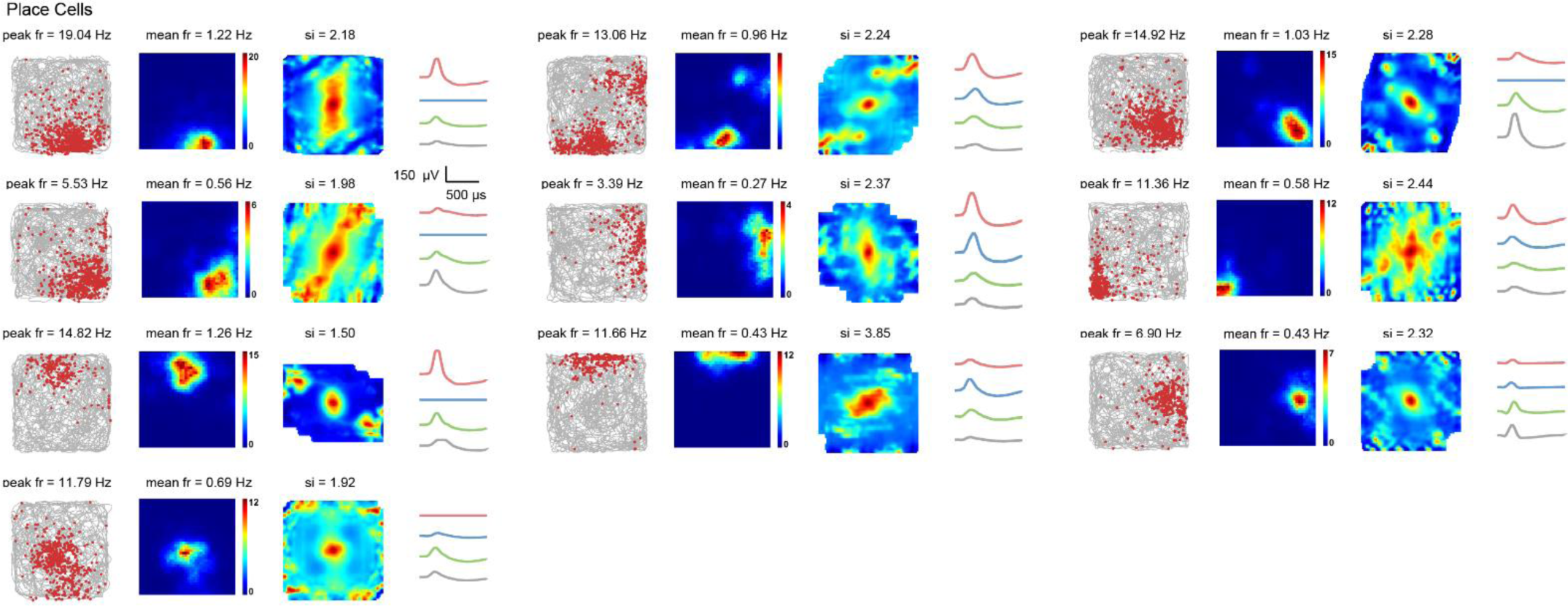
Entire samples of V2 place cells. Trajectory (grey line) with superimposed spike locations (red dots) (first column); spatial firing rate maps (second column) and autocorrelation diagrams (third column). Firing rate was color-coded with dark blue (red) indicating minimal (maximal) firing rate. The scale of the autocorrelation maps was twice that of spatial firing rate maps. Peak firing rate, mean firing rate, and spatial information (SI) for each representative V2 place cell are labelled at the top of panels. Fourth column: Average spike waveforms on four electrodes of the same tetrode. Scale bar, 150 µV and 500 µs.

**Fig. S8.**
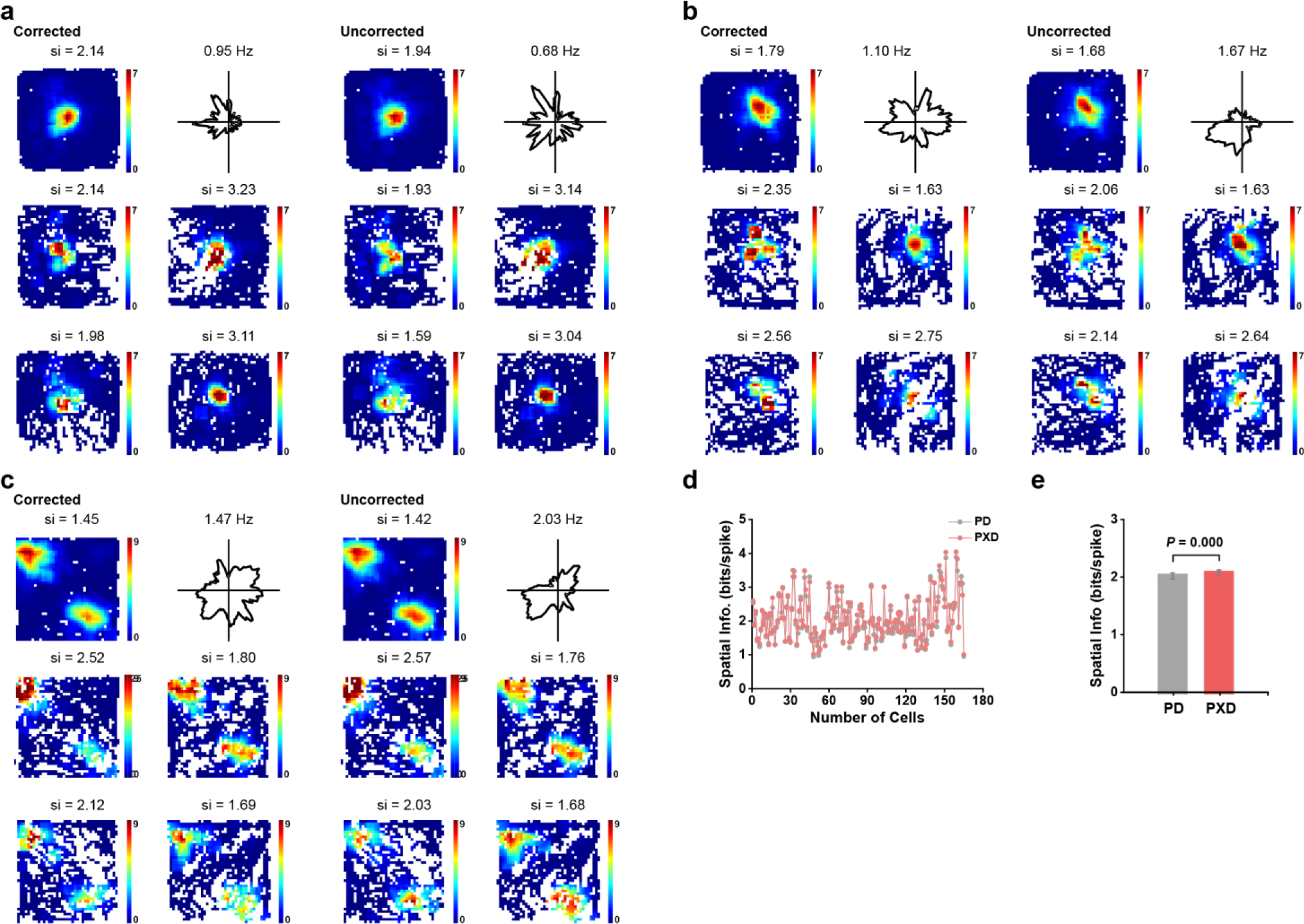
Maximum likelihood estimation of the tuning property of V2 place cells. **a-c**, Tuning properties of representative V2 place cells in **Fig.1b** using maximum likelihood estimation. First two columns, corrected responses with maximum likelihood model; Last two columns, uncorrected responses. Top two panels show rate map and head direction tuning. Data from four head direction ranges (0°-90°, 90°-180°, 180°-270°, 270°- 360°) are shown in the middle and bottom four panels. Note the similar spatial rate maps in four head direction ranges. **d**, Distribution of spatial information for all recorded V2 place cells (PD, uncorrected; PXD corrected). **e**, The increase in spatial information after using maximum likelihood estimation algorithm compared to the uncorrected spatial information (*n* = 165, two-tailed paired *t*-test, *P* = 0.000).

**Fig. S9.**
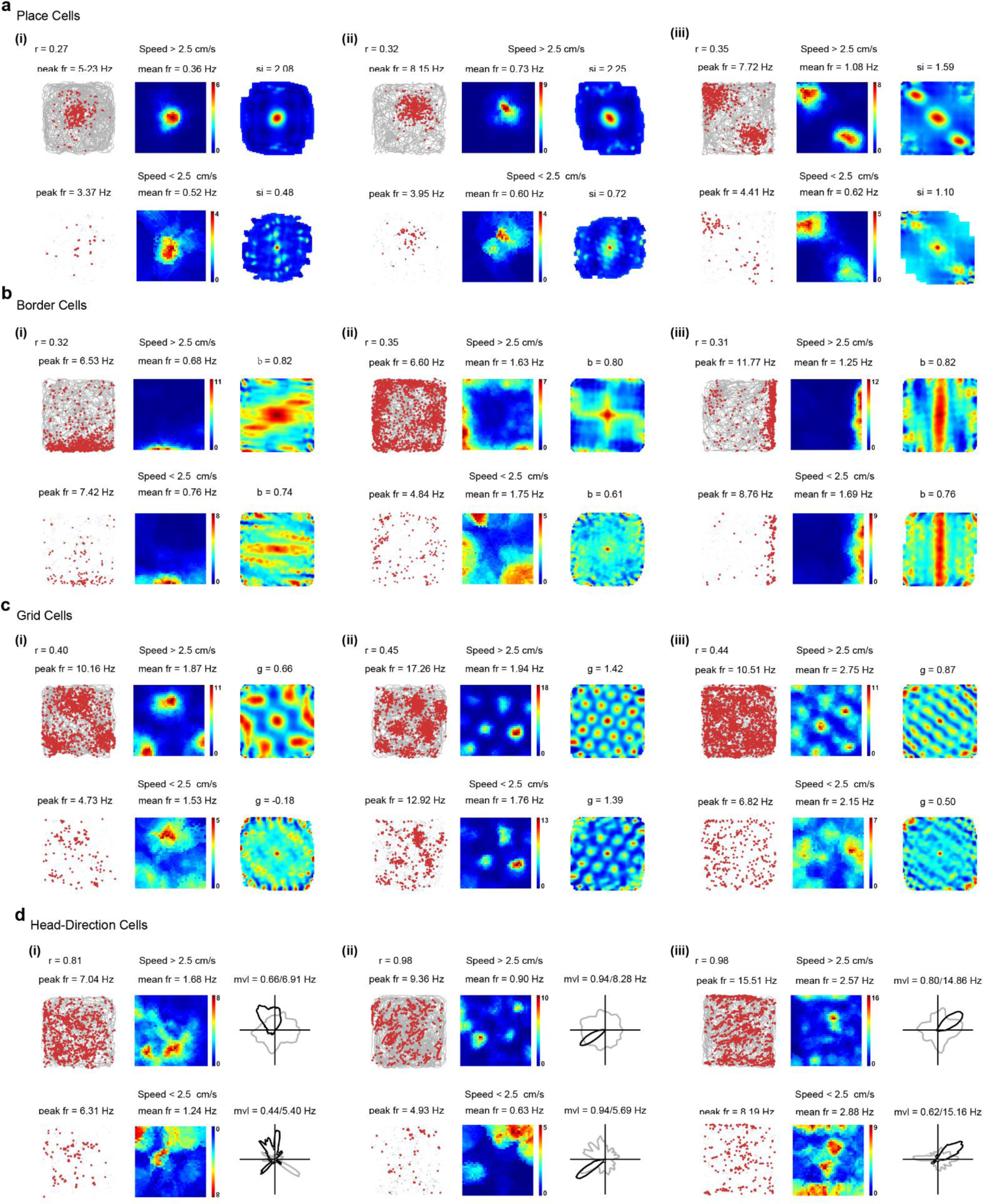
Comparison of spatial responses of V2 spatially tuned cells between running and slow ent. a-d, Tuning properties of representative V2 place cells in Fig. lb, V2 border cells in Fig. 2a, V2 lls in Fig.3a, and V2 head-direction cells in Fig. 4a during running (running speed > 2.5 cm/s) and unning or immobility (running speed < 2.5 crn/s). Trajectory (grey line) with superimposed spike ns (red dots) (left column); spatial firing rate maps (middle column) and autocorrelation diagrams (right). Firing rate was color-coded with dark blue (red) indicating minimal (maximum) firing. The scale of ocorrelation maps was twice that of spatial firing rate maps. Peak firing rate, mean firing rate, spatial ation, border score or gridness score are shown. Third column: For head-direction cells, head direction curves (black) plotted against dwell-time polar plot (grey), and the numbers represent the angular peak d mean vector length; for other cell types, autocorrelation maps were plot.

**Fig. S10.**
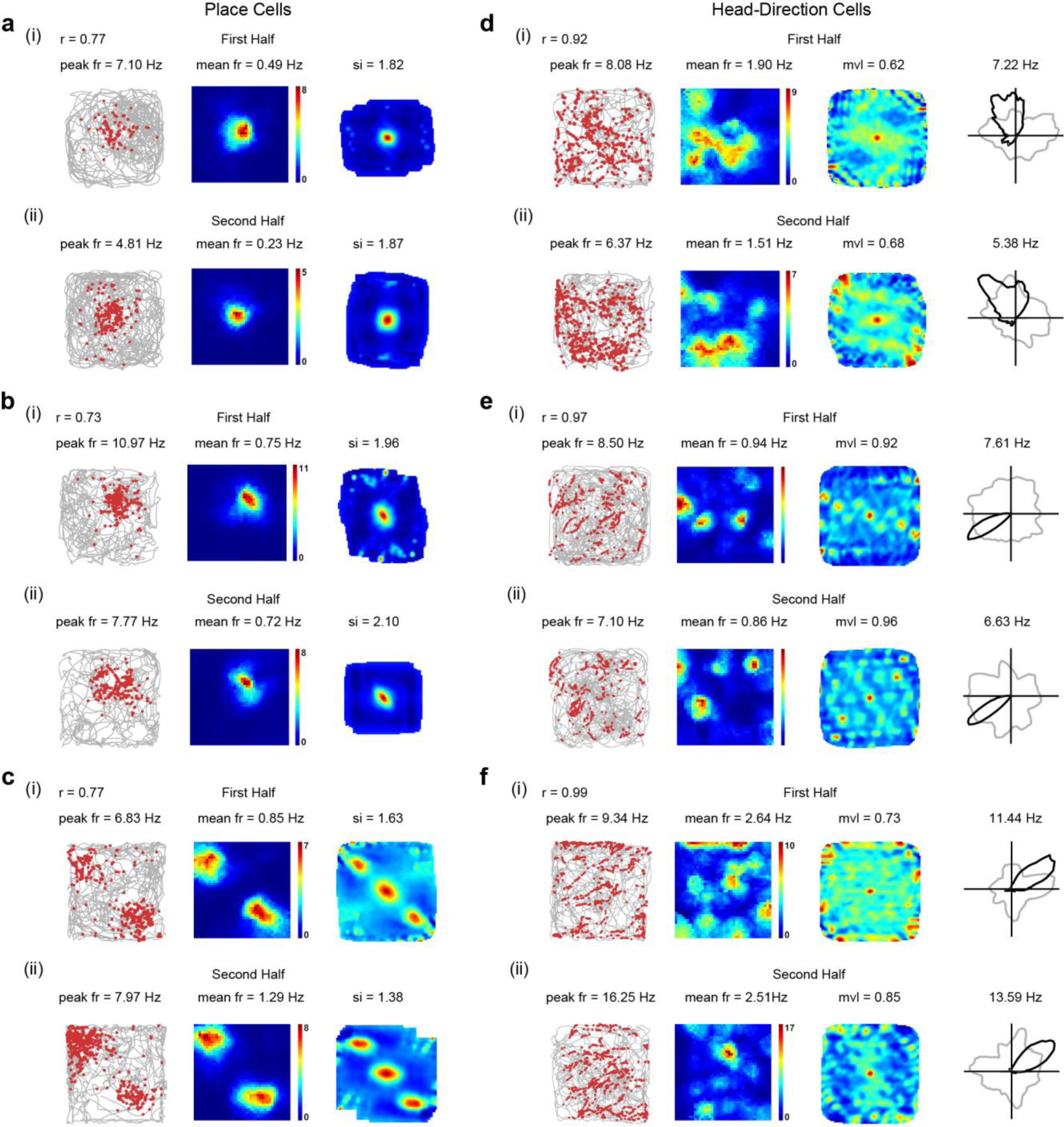
Stable tuning of V2 place cells and V2 head-direction cells. **a-c**, Spatial responses of the three representative V2 place cells in Fig. 1b for the first half and second half of recording sessions. V2 place cells exhibit stable spatial firing patterns. Correlation coefficient (r) of spatial maps between the first half and second half of the recording session are shown for three V2 place cells. **d-f**, Spatial responses of the three representative V2 head-direction cells in Fig. 4a for the first half and second half of recording sessions. V2 head-direction cells exhibit stable preferred firing directions. Correlation coefficient of the angular map between the first half and second half of recording sessions are shown for three V2 head-direction cells.

**Supplementary Fig. S11.**
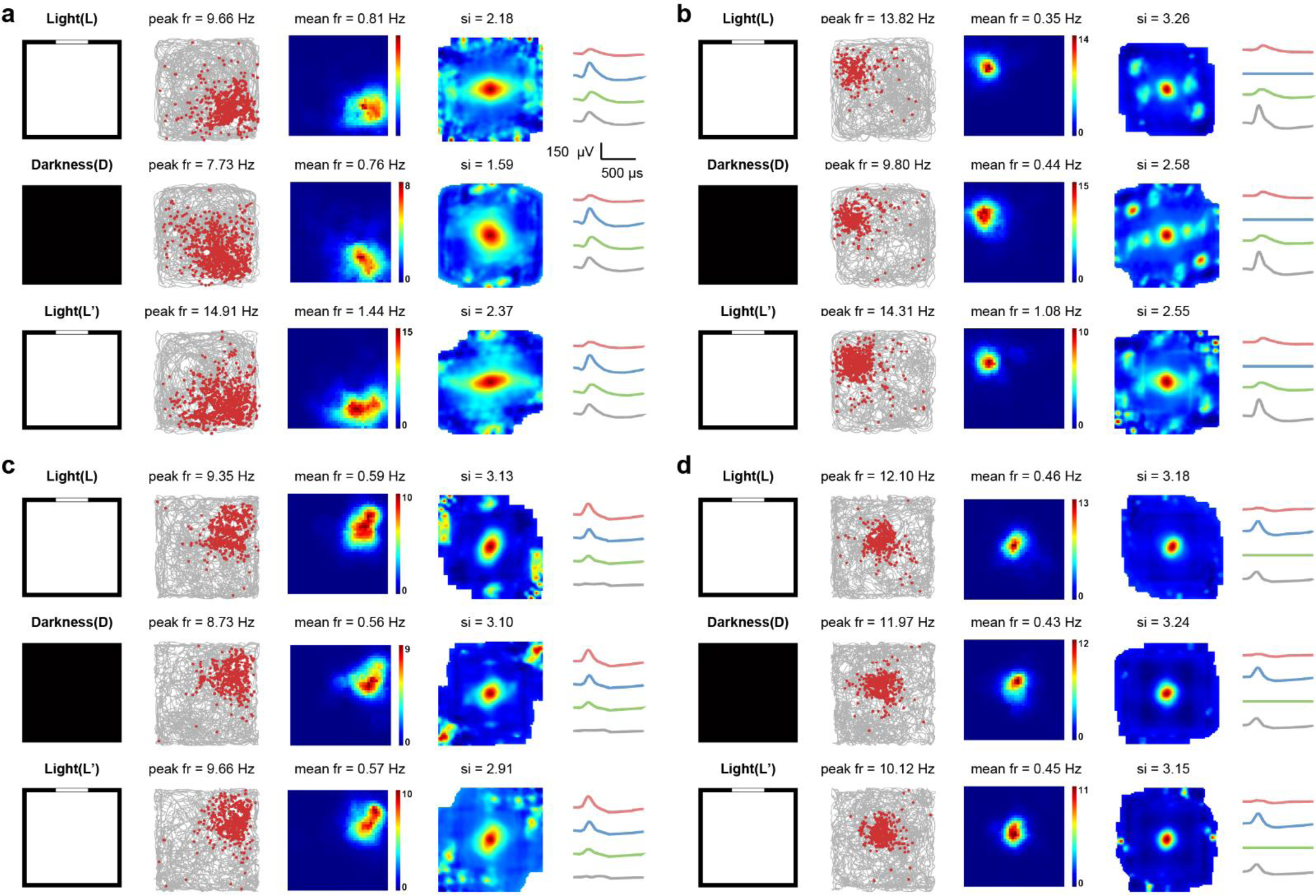
Preserved spatial selectivity of V2 place cells in the darkness. **a-d**, Responses of four representative V2 place cells in the darkness. Responses of V2 place cells in the L-D-L’ conditions. Symbols and notations are similar as before.

**Fig. S12.**
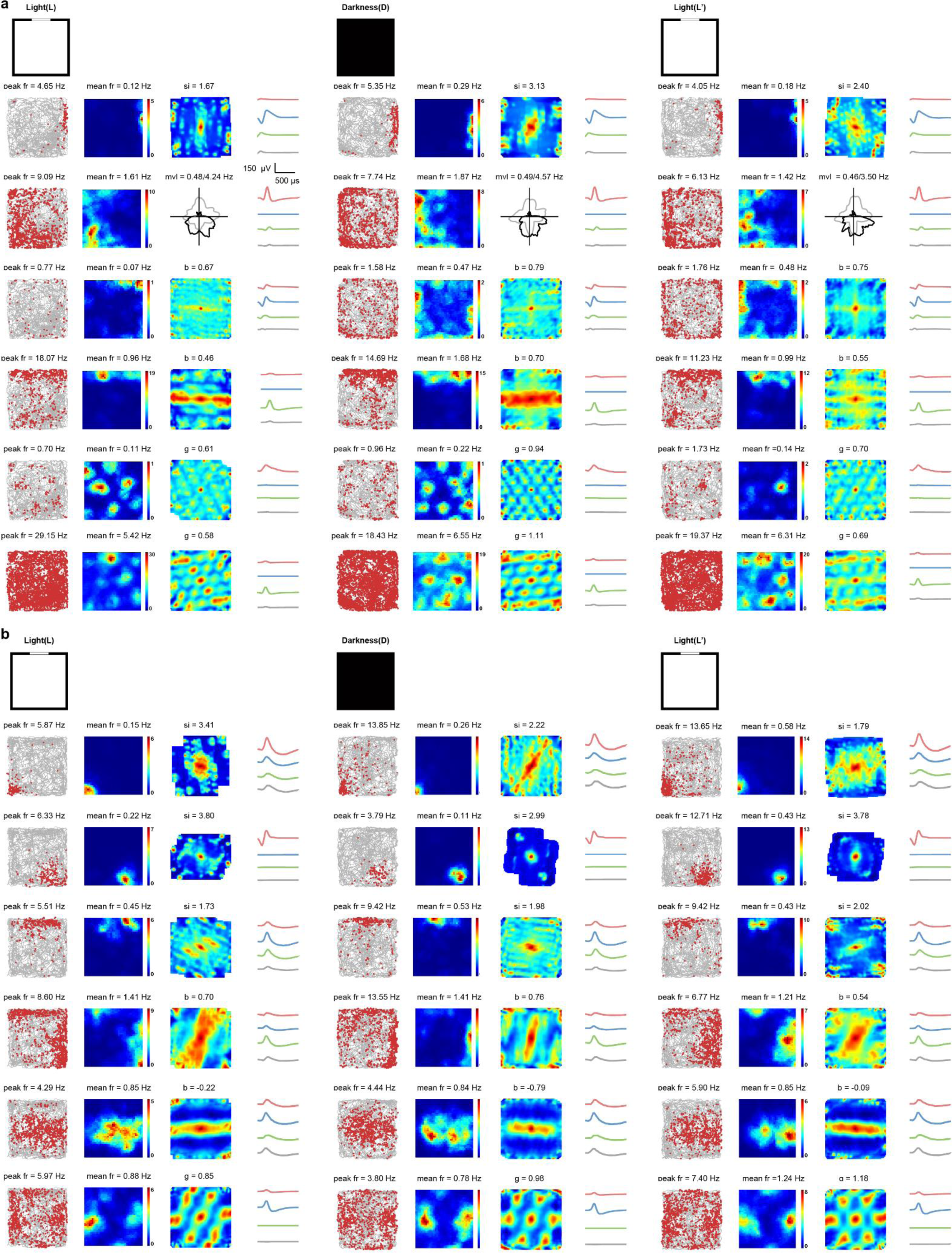
Persistence of spatial responses of simultaneously recorded multiple types ofV2 spatially tuned darkness. **a,** Spatial firing of six simultaneously recorded V2 units (from top to bottom: place cell, irection cell, border cell, border cell, grid cell and grid cell) in the L-D-L’ conditions. Symbols and ns are similar as before. **b,** Same as **a** but for a different set of six simultaneously recorded V2 units op to bottom: place cell, place cell, place cell, border cell, band cell and grid cell).

**Fig. S13.**
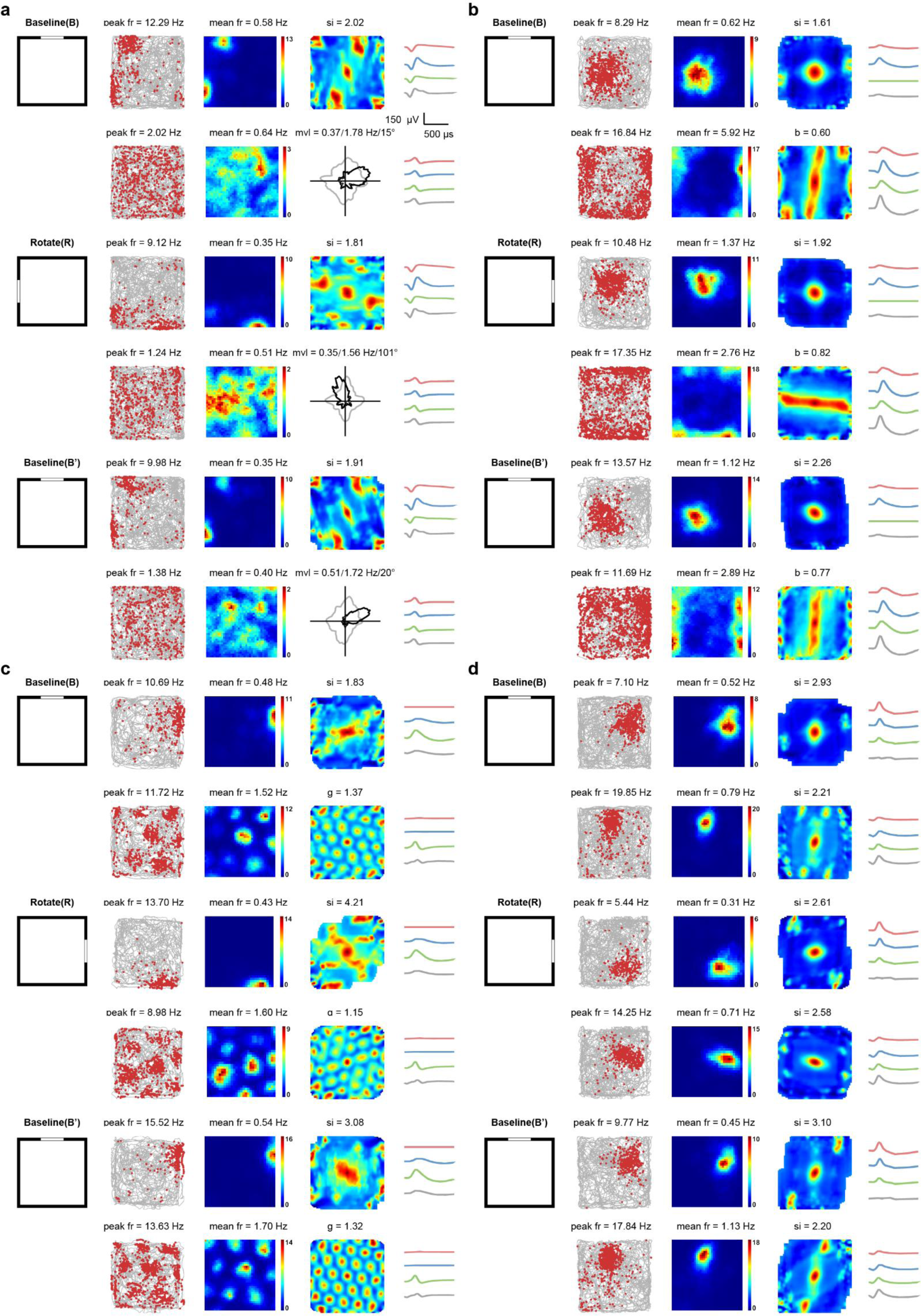
Visual landmark control of V2 place cells and simultaneously recorded other spatially tuned cells. **a**, One V2 place cell with one simultaneously recorded V2 head-direction cell in response to visual landmark manipulation. **b**, One V2 place cell with one simultaneously recorded V2 border cell in response to visual landmark rotation. **c**, One V2 place cell with one simultaneously recorded V2 grid cell in response to visual cue manipulation. **d**, Two simultaneously recorded V2 place cells in response to visual landmark manipulation. Top to bottom panels: responses of V2 place cells and simultaneously recorded other V2 spatially tuned cells in baseline, clockwise or counterclockwise 90°degree rotation of visual cue and back to baseline. Symbols and notations are similar as before.

**Fig. S14.**
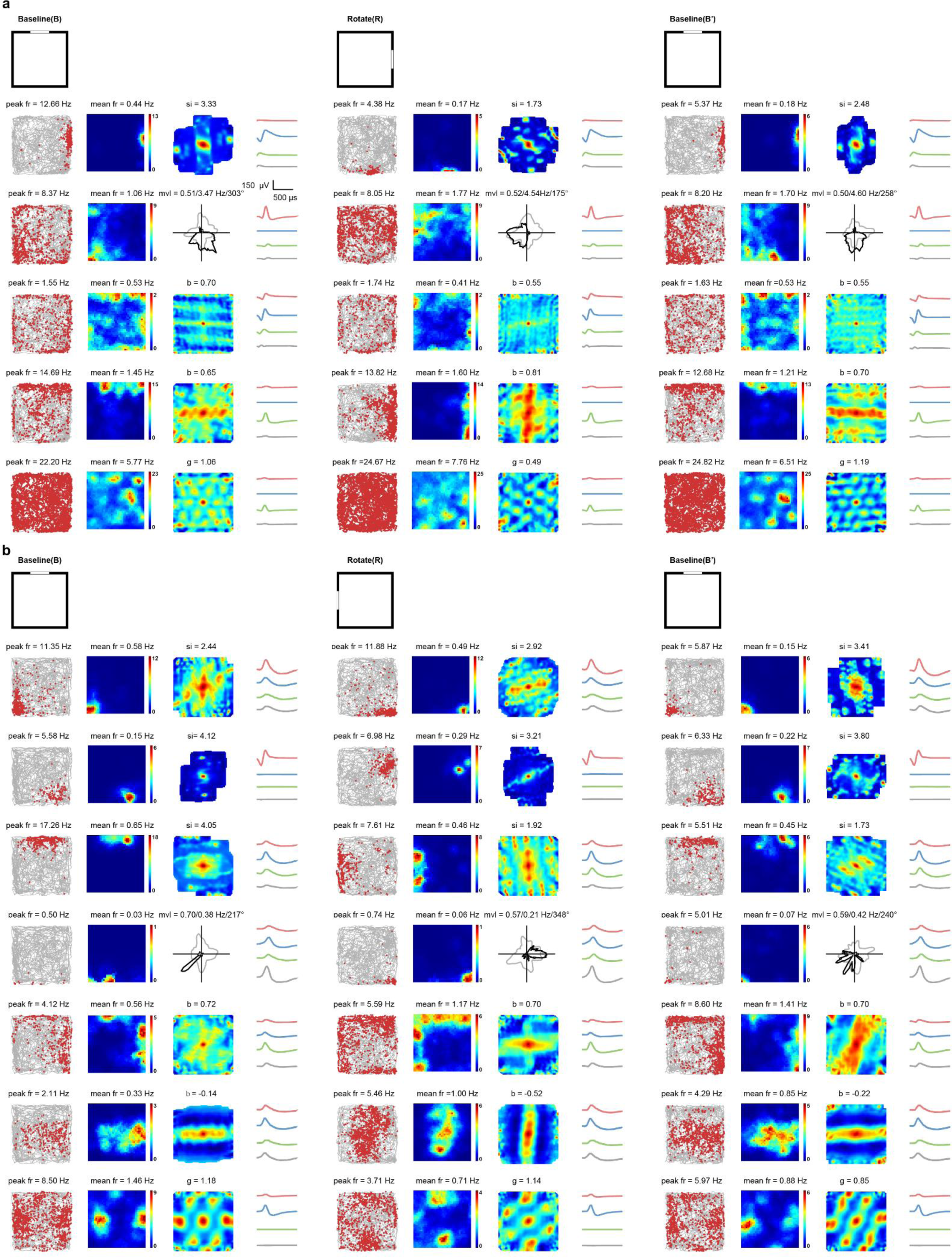
Visual landmark control of simultaneously recorded multiple types ofV2 spatially tuned cells. **a**, Spatial firing of five simultaneously recorded V2 units (from top to bottom: place cell, head-direction cell, cell, border cell and grid cell) in response to clockwise 90° visual cue rotation. Symbols and notations ilar as before. **b,** Spatial firing of seven simultaneously recorded V2 units in another independent ng session (from top to bottom: place cell, place cell, place cell, head-direction cell, border cell, band d grid cell) in response to counterclockwise 90° visual cue rotation.

**Supplementary Fig. S15.**
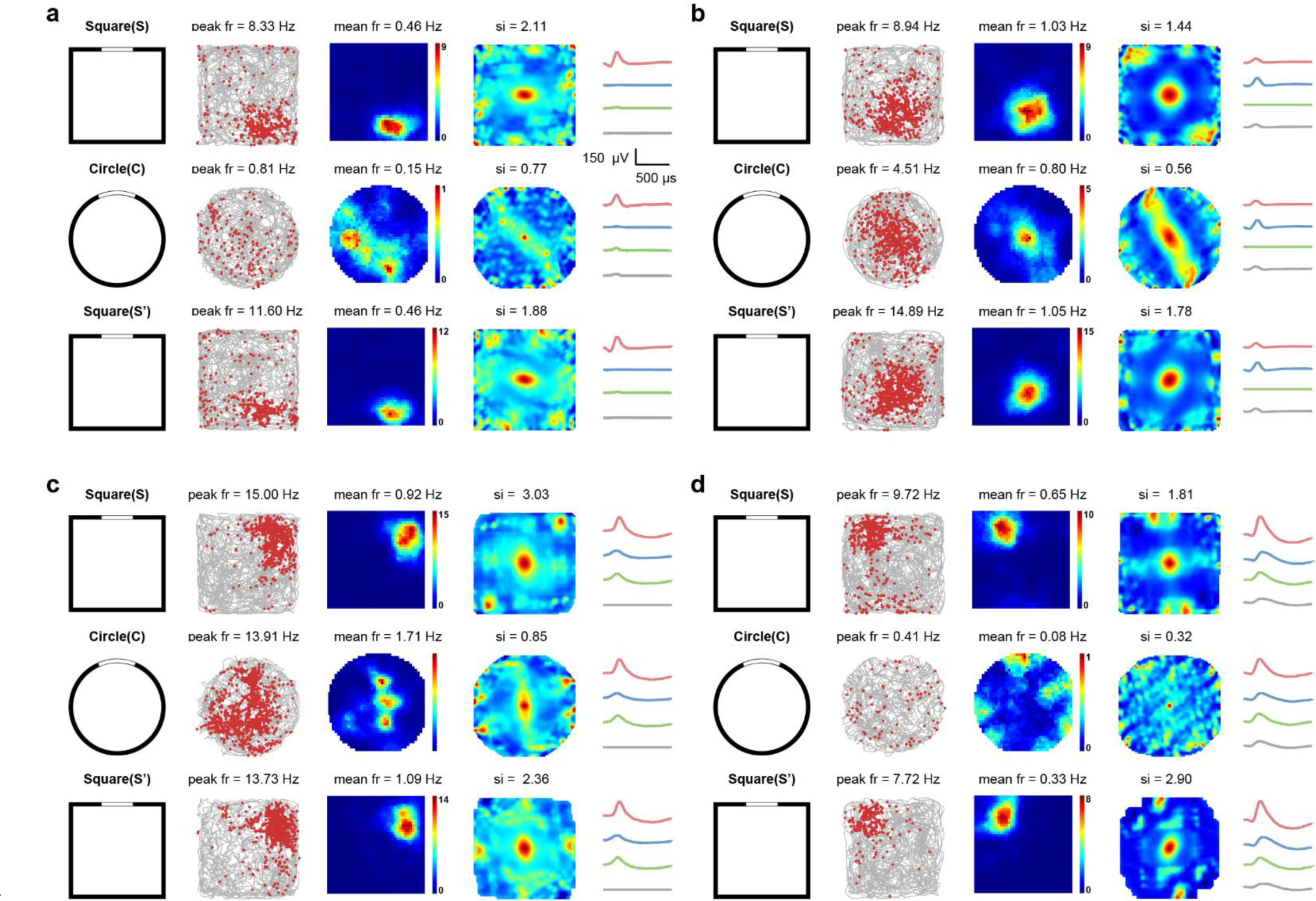
Remapping of V2 place cells in different geometric environments. **a-d**, Spatial responses of four representative V2 place cells showed remapping in response to different environmental shapes. Upper to bottom panels, responses of the place cells in the square enclosure (upper panel), circle (middle panel) and back to the square enclosure (lower panel). Symbols and notations are similar as before.

**Fig. S16.**
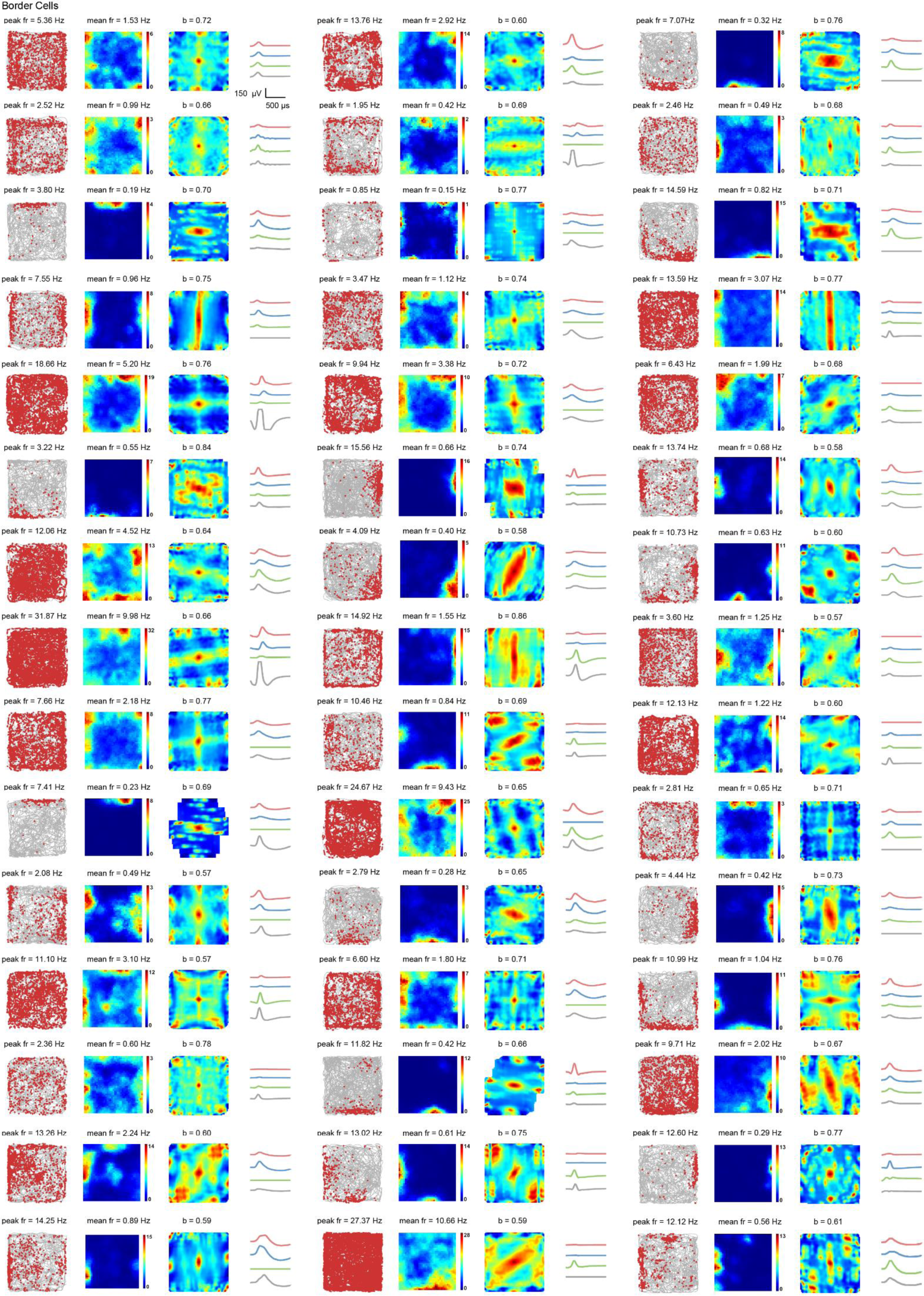
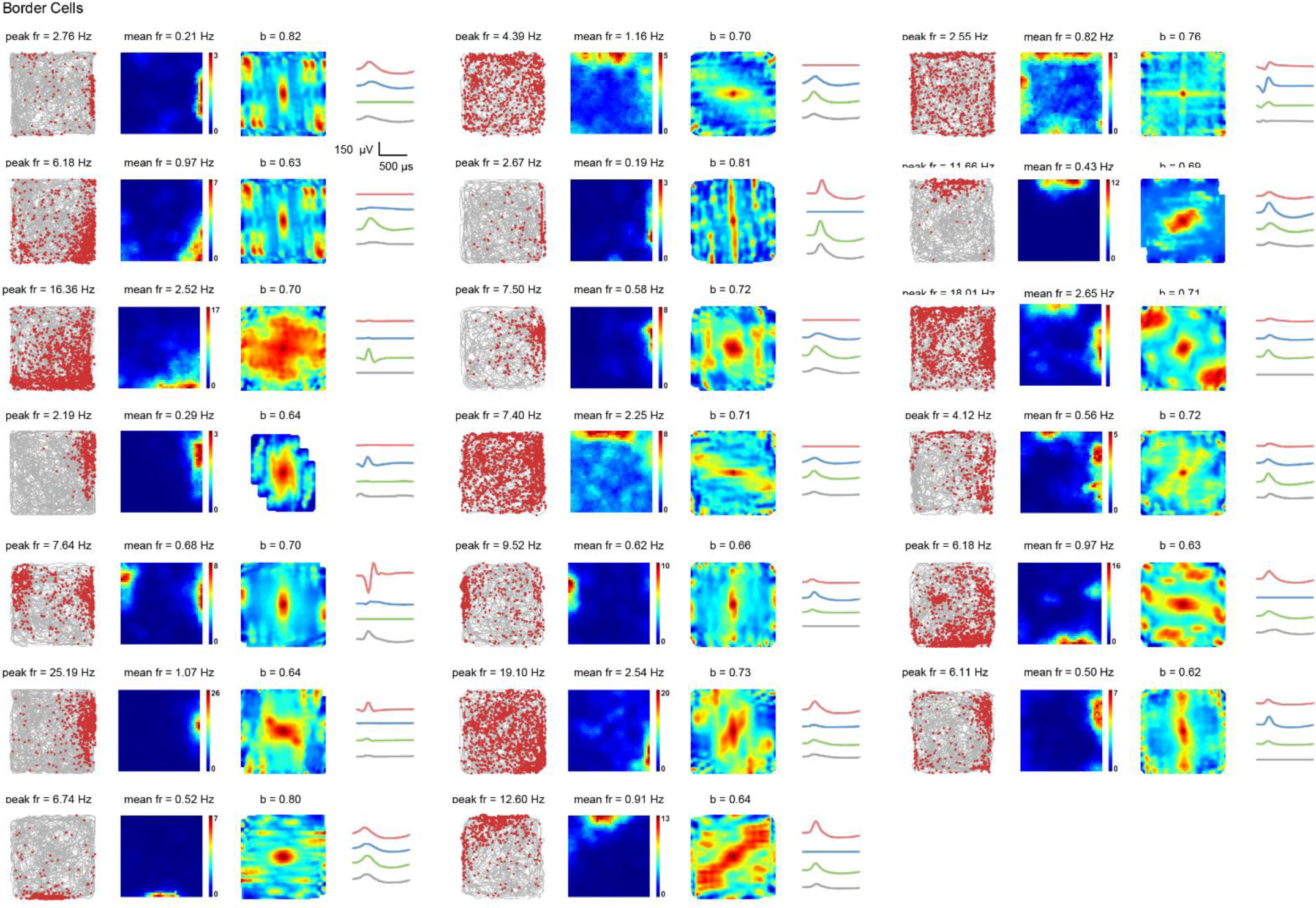
Entire samples of V2 border cells. Trajectory (grey line) with superimposed spike locations (red dots) (first column); spatial firing rate maps (second column) and autocorrelation diagrams (third column). Firing rate was color-coded with dark blue (red) indicating minimal (maximum) firing rate. The scale of the autocorrelation maps was twice that of the spatial firing rate maps. Symbols and notations are similar as before.

**Fig. S17.**
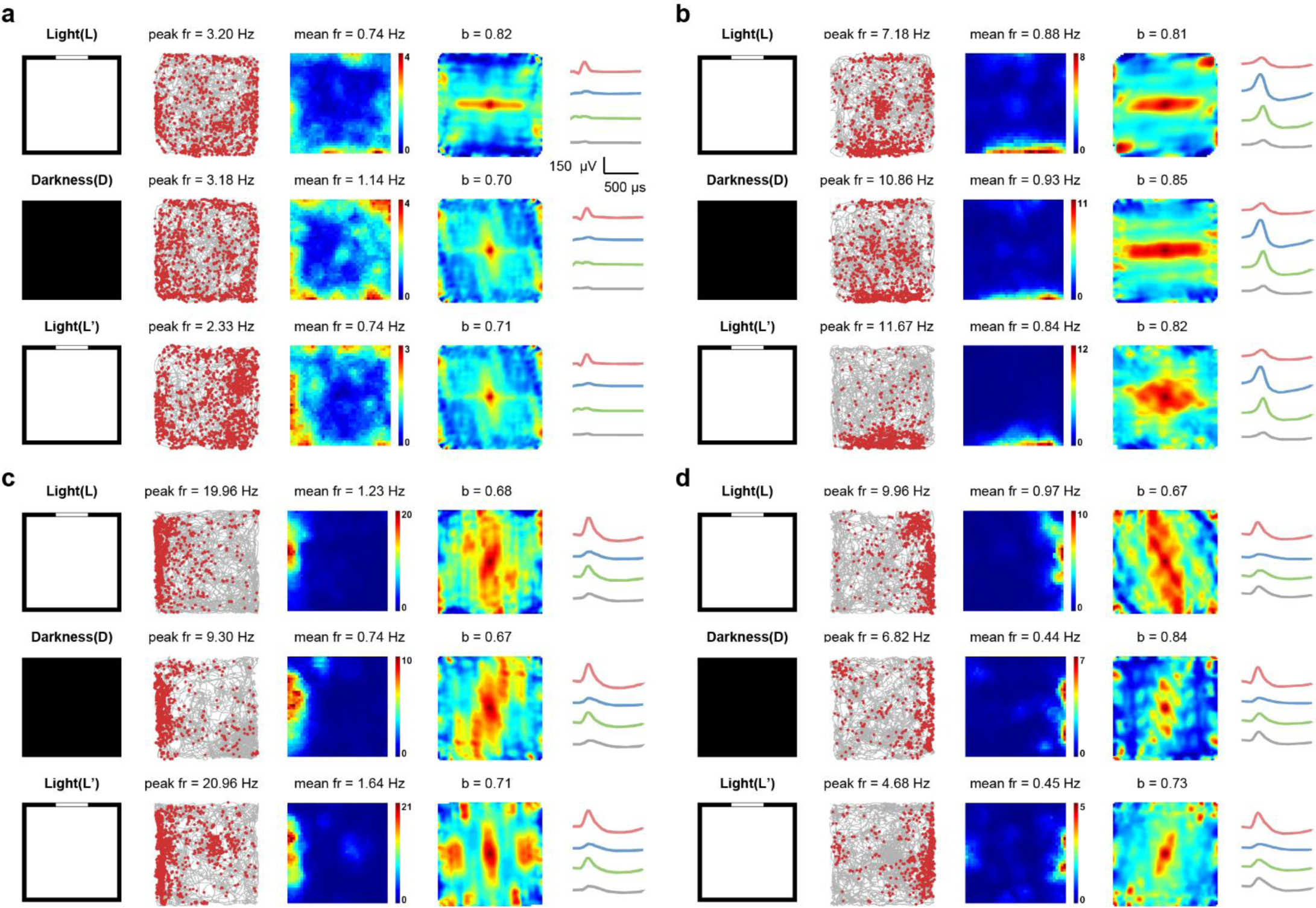
V2 border cells preserved spatial firing patterns in the darkness. **a-d**, Responses of representative V2 border cells in the darkness. Symbols and notations are similar as before.

**Fig. S18.**
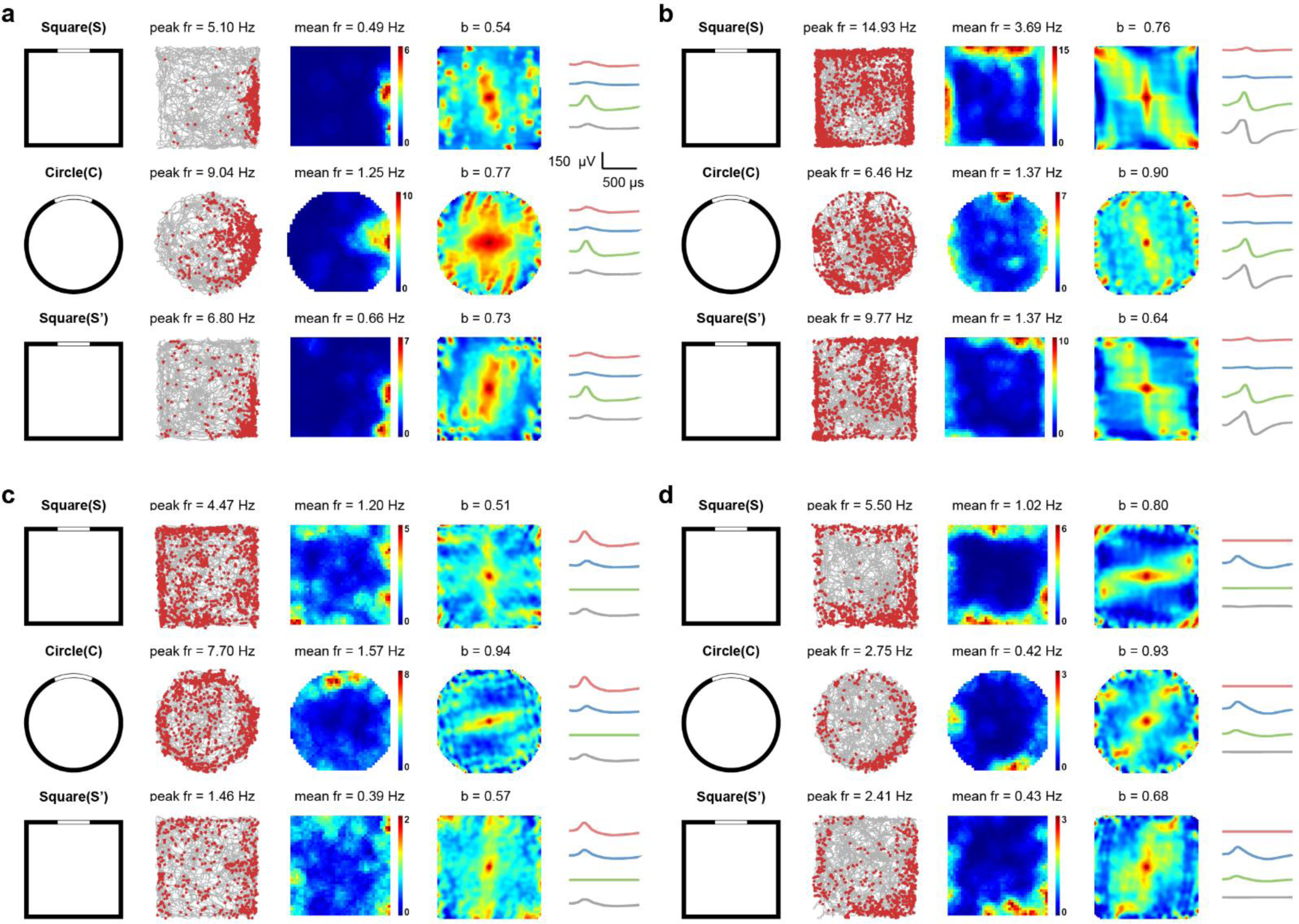
Preserved spatial selectivity of V2 border cells in different shapes of environments. **a-d**, Responses of four representative V2 border cells in navigation environments with different shapes. Symbols and notations are similar as before.

**Fig. S19.**
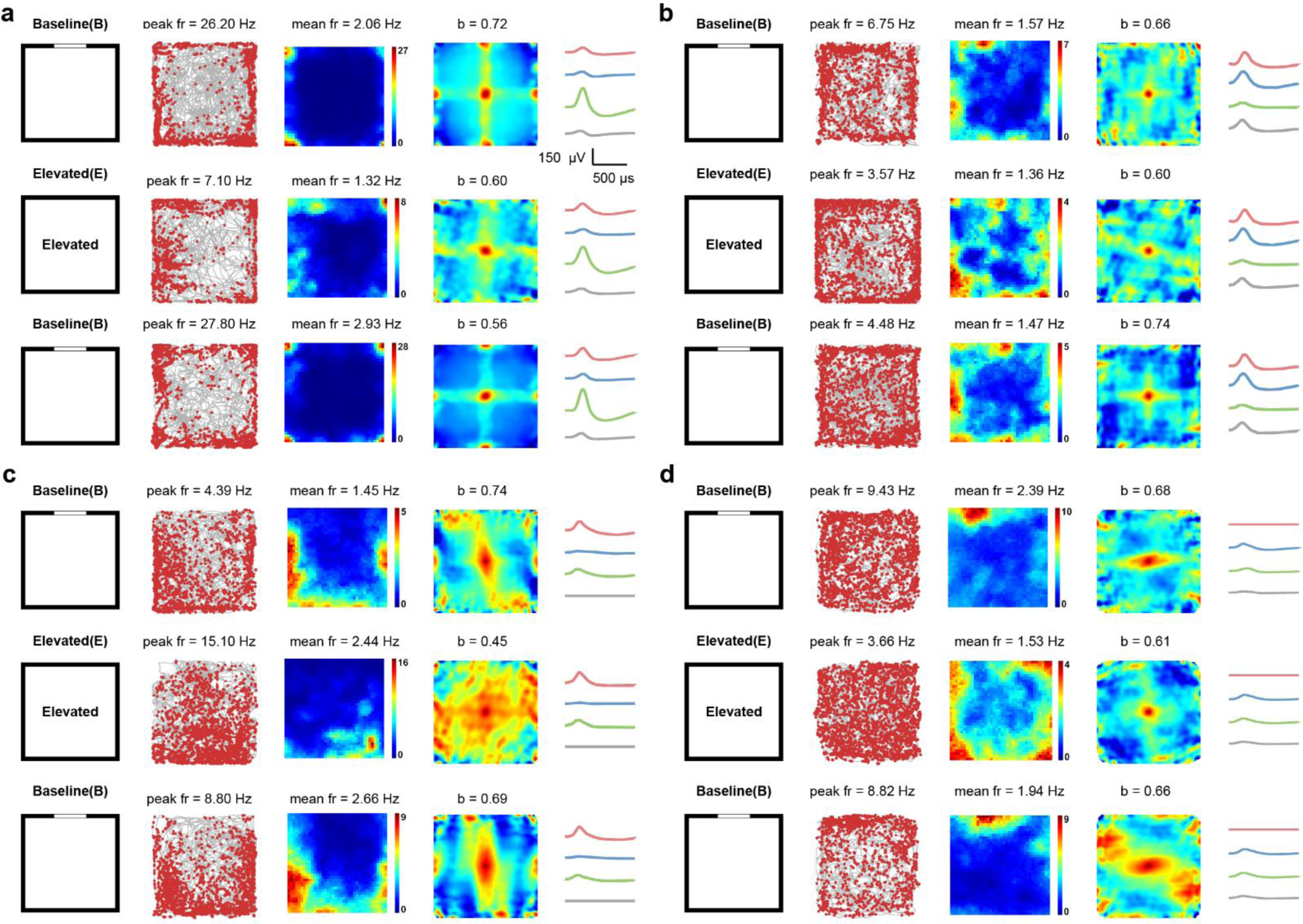
Spatial responses of V2 border cells in the elevated platform without walls. **a-d**, Responses of representative V2 border cells in the elevated platform without walls. First column: schematic of experimental paradigms. Symbols and notations are similar as before.

**Fig. S20.**
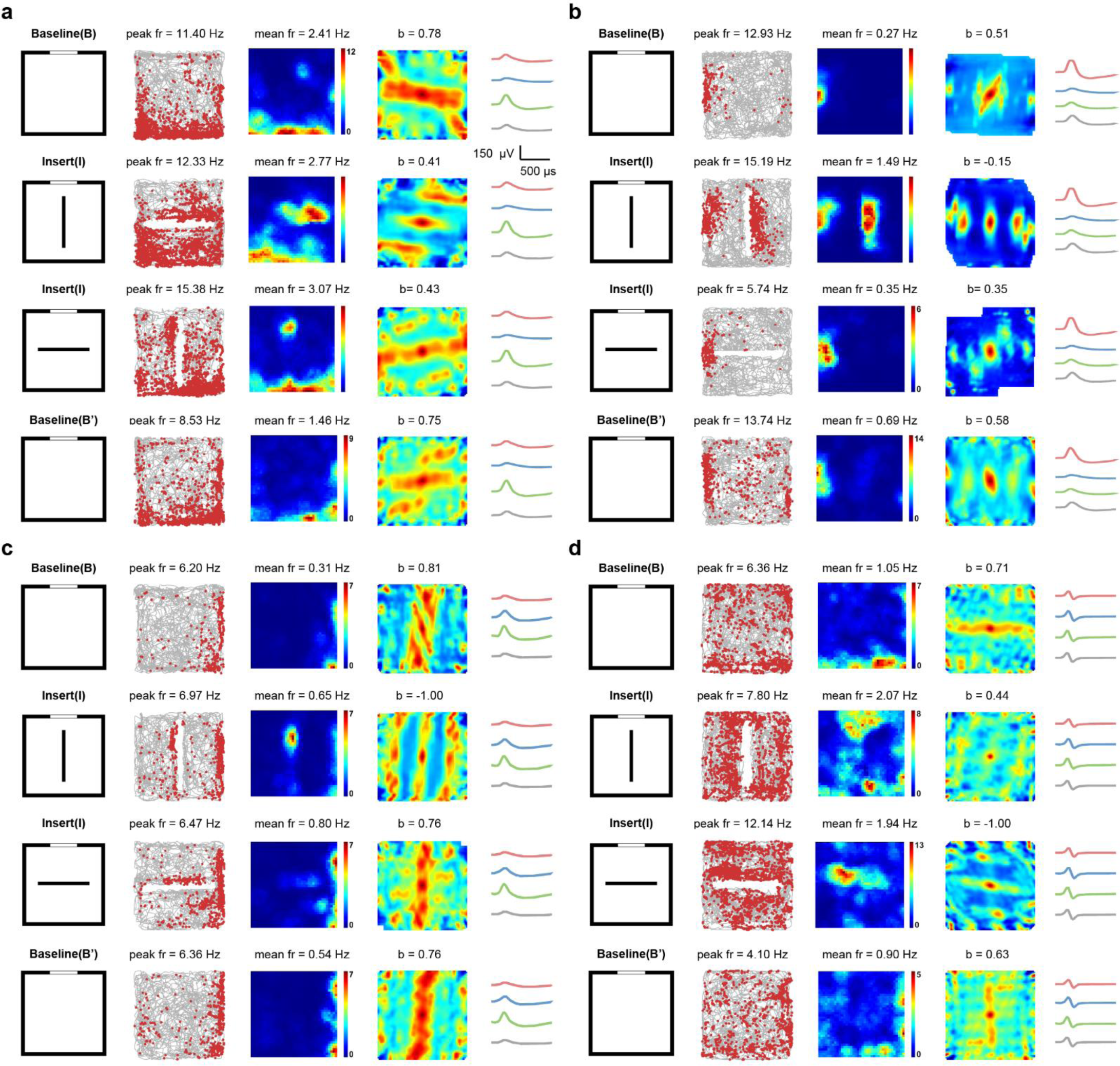
Spatial responses of V2 border cells to newly inserted walls. **a-d**, Responses of four representative V2 border cells to the external insert. Left column: Schematic of experimental paradigms. Symbols and notations are similar as before.

**Fig. S21.**
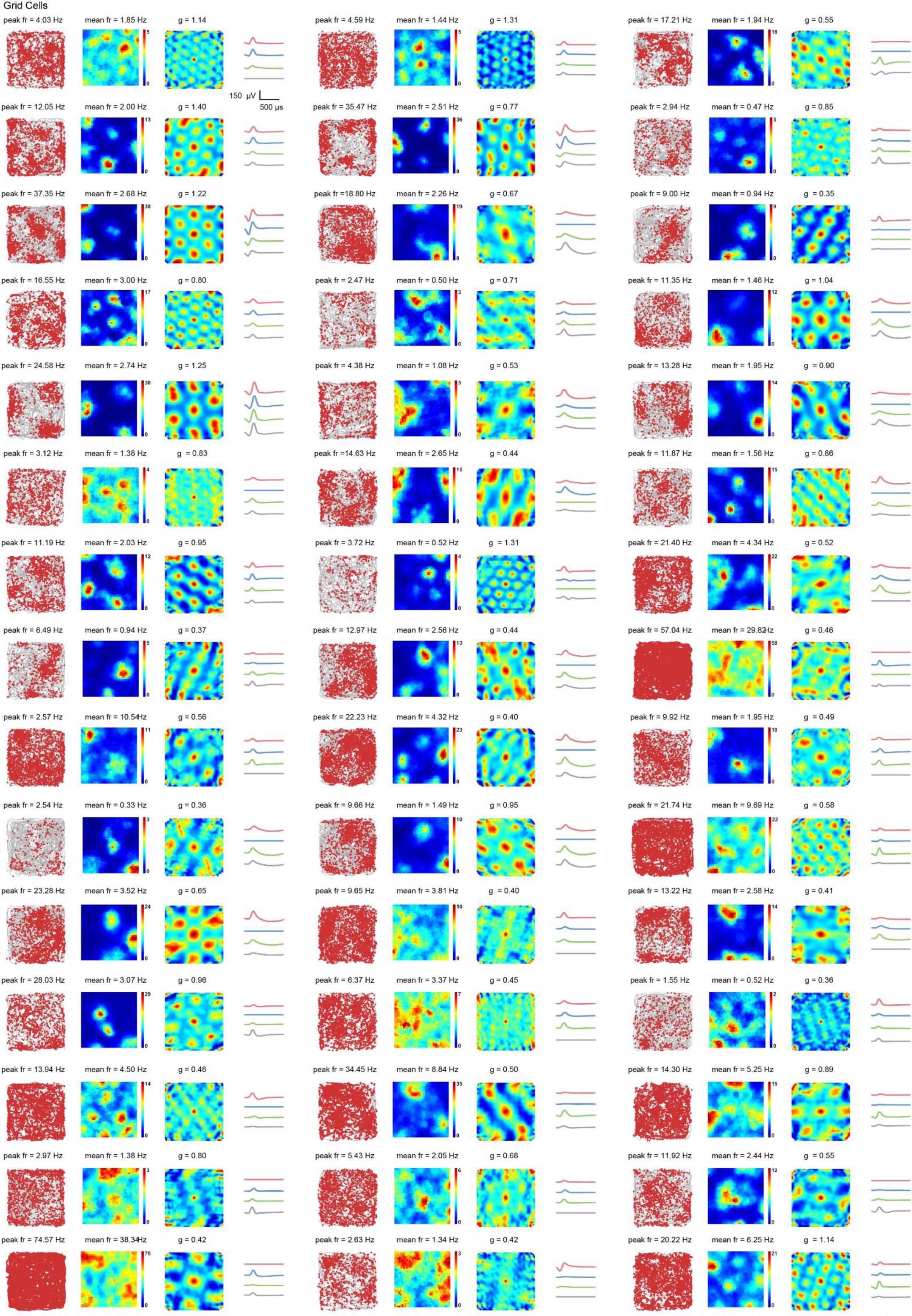
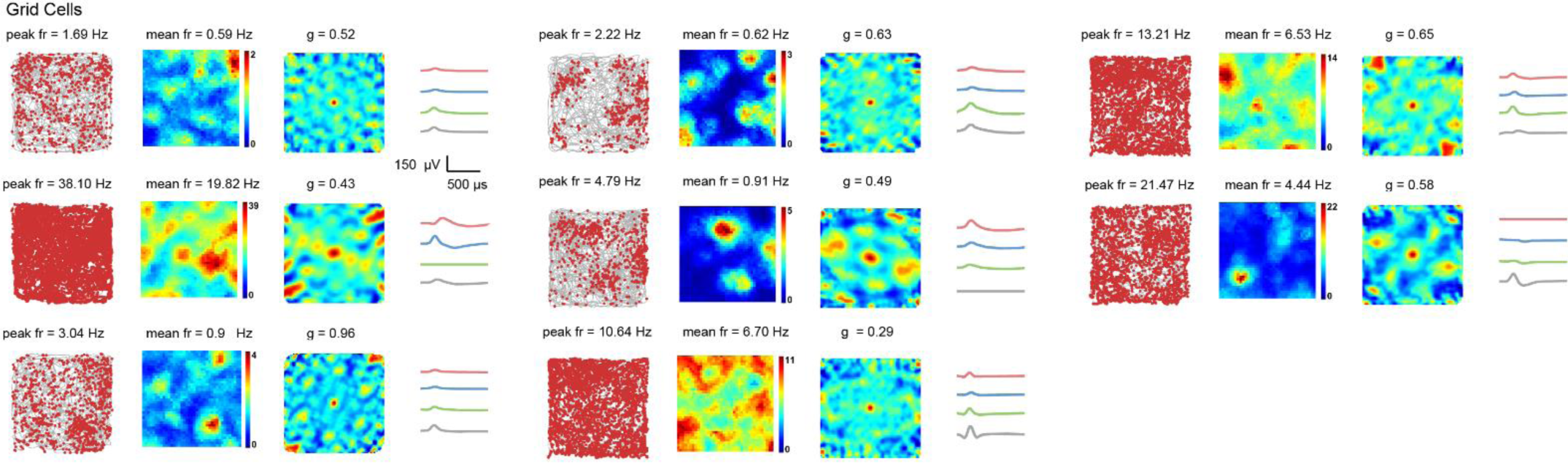
All identified samples of V2 grid cells. Symbols and notations are similar as before.

**Fig. S22.**
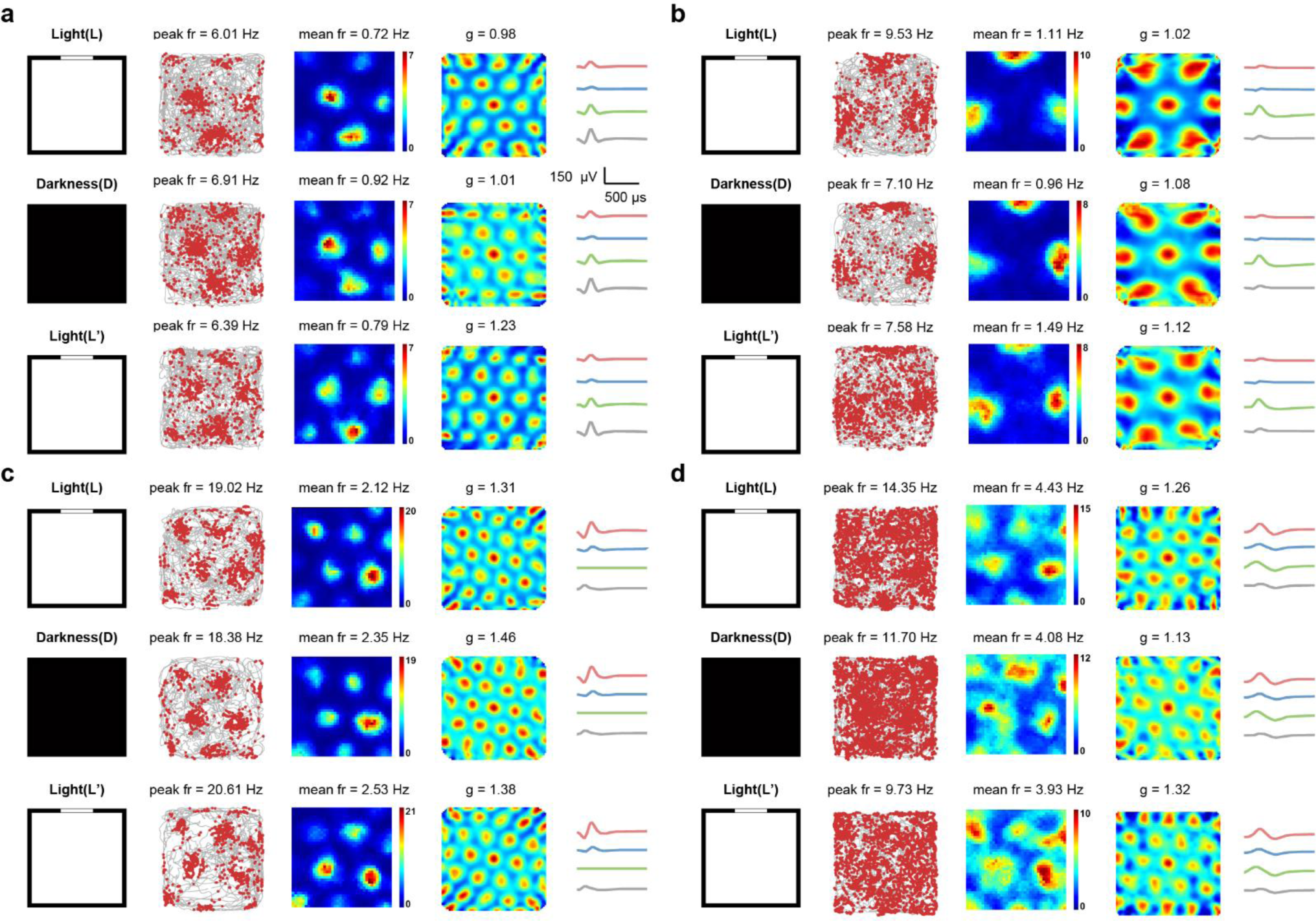
Stable hexagonal spatial firing patterns of V2 grid cells in the darkness. **a-d**, Responses of four representative V2 grid cells in the darkness. Schematic of the L-D-L’ experimental paradigm. Symbols and notations are similar as before.

**Fig. S23.**
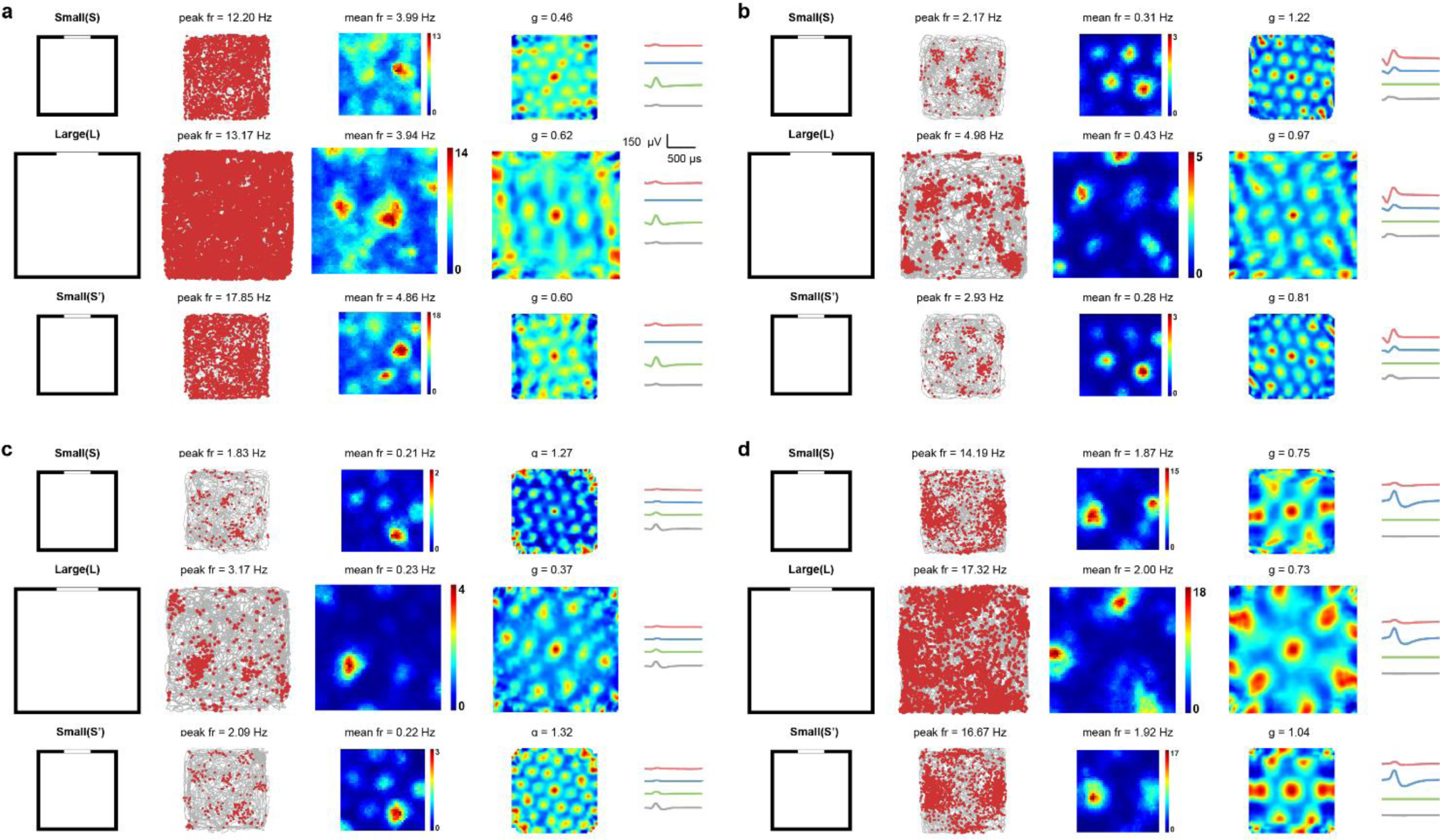
Persistence of grid firing patterns of V2 gird cells in larger environments. **a-d**, Responses of representative V2 grid cells in running environments with different sizes. Left column: schematic of experimental paradigms during switch of environment: 1 ×1 m^2^ (S), 1.5 ×1.5 m^2^ (L) and 1 ×1 m^2^ (S’). Symbols and notations are similar as before.

**Fig. S24.**
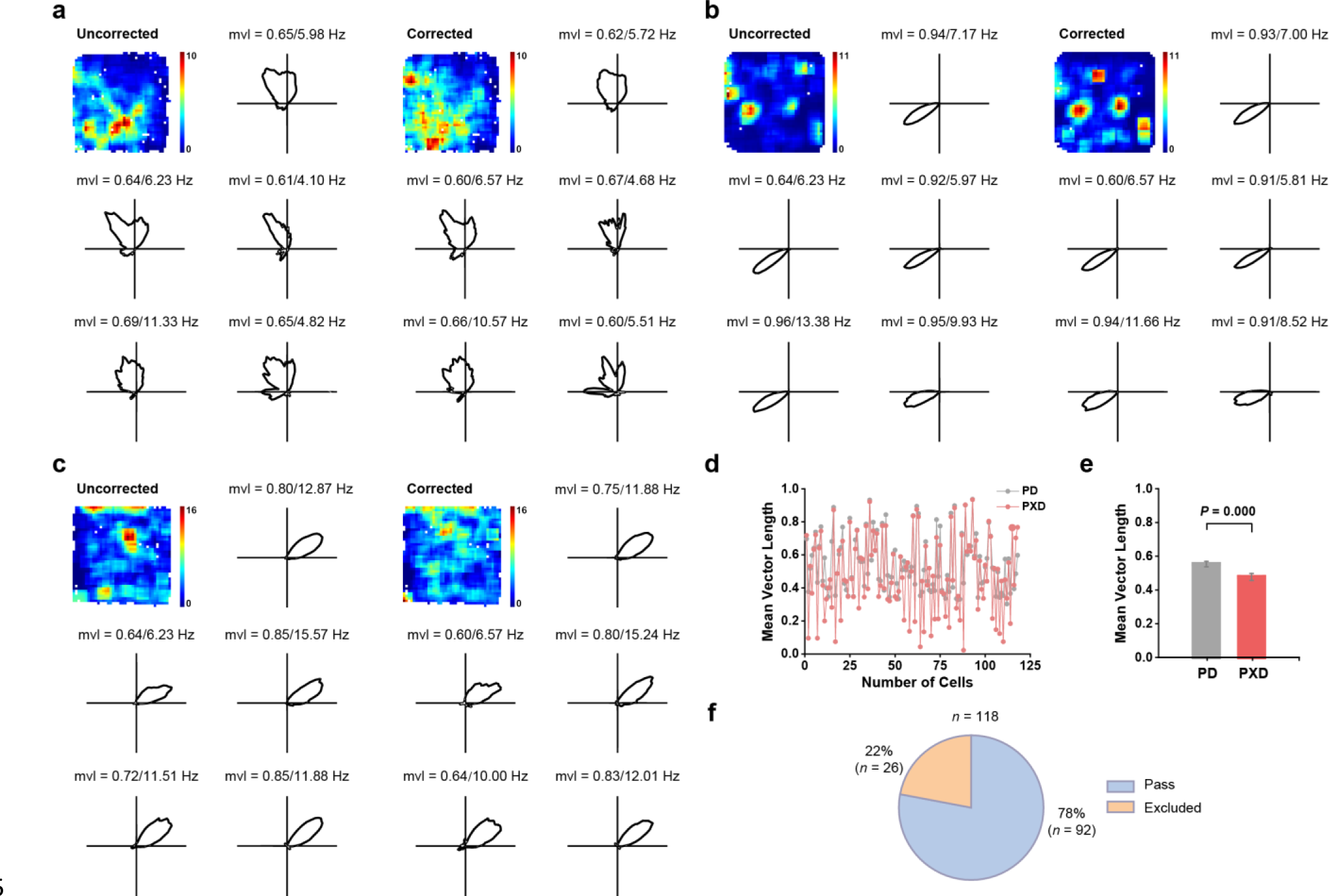
Maximum likelihood estimation of the tuning property of V2 head-direction cells. **a-c**, Tuning properties of representative V2 head-direction cells in Fig. 4a using maximum likelihood correction. First two columns: corrected responses; last two columns: uncorrected responses. Top row shows the rate map and head direction tuning. Bottom two rows show the data from the four quadrant. Note that the preferred directions were similar in the four quadrants of the environment. **d**, Distribution of mean vector length for all identified V2 head-direction cells that passed the threshold criterion (PD, uncorrected; PXD corrected). **e**, Mean vector length after maximum likelihood estimation was significantly reduced compared to the uncorrected mean vector length (*n* = 116, two-tailed paired *t*- test, *P* = 0.000). **f**, Venn diagram showing the proportion of V2 head-direction cells passing (92 out of 118, 78%) or excluded (26 out of 118, 22%) by the maximum likelihood estimation algorithm.

**Fig. S25.**
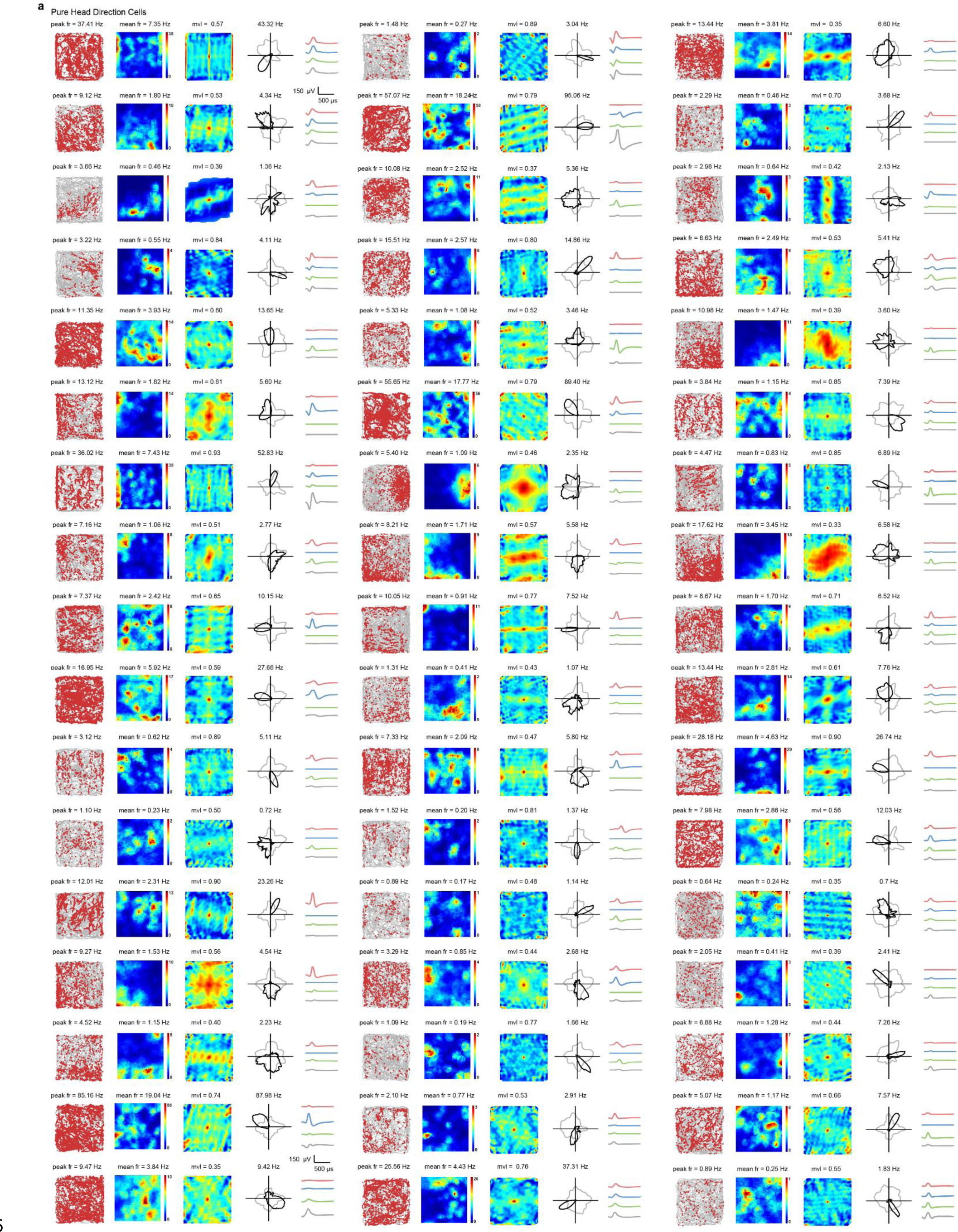
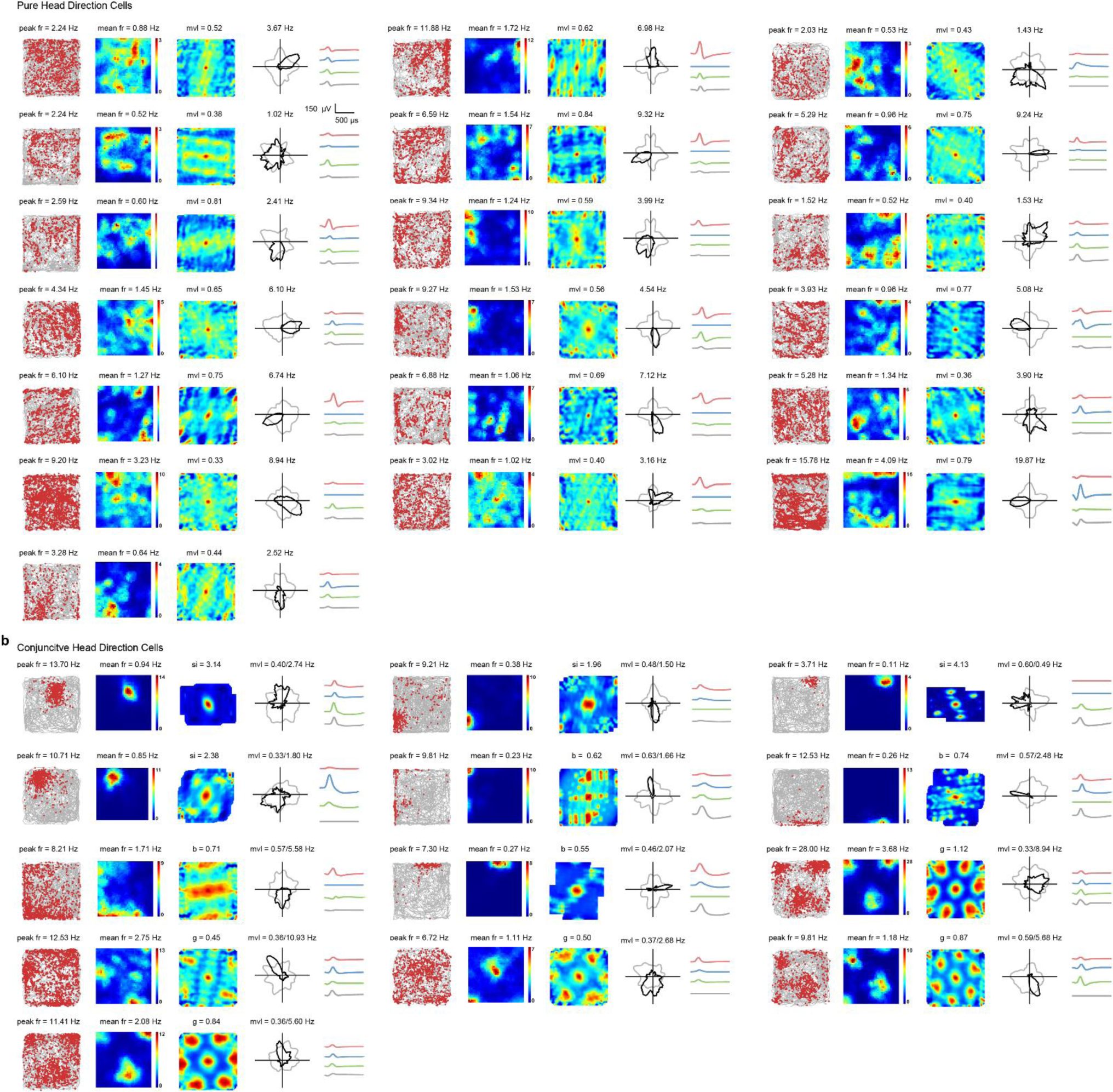
All identified pure V2 head-direction cells and conjunctive V2 head-direction cells. **a**, Pure V2 head-directions cells. **b**, Conjunctive V2 head-direction cells. Symbols and notations are similar as before.

**Fig. S26.**
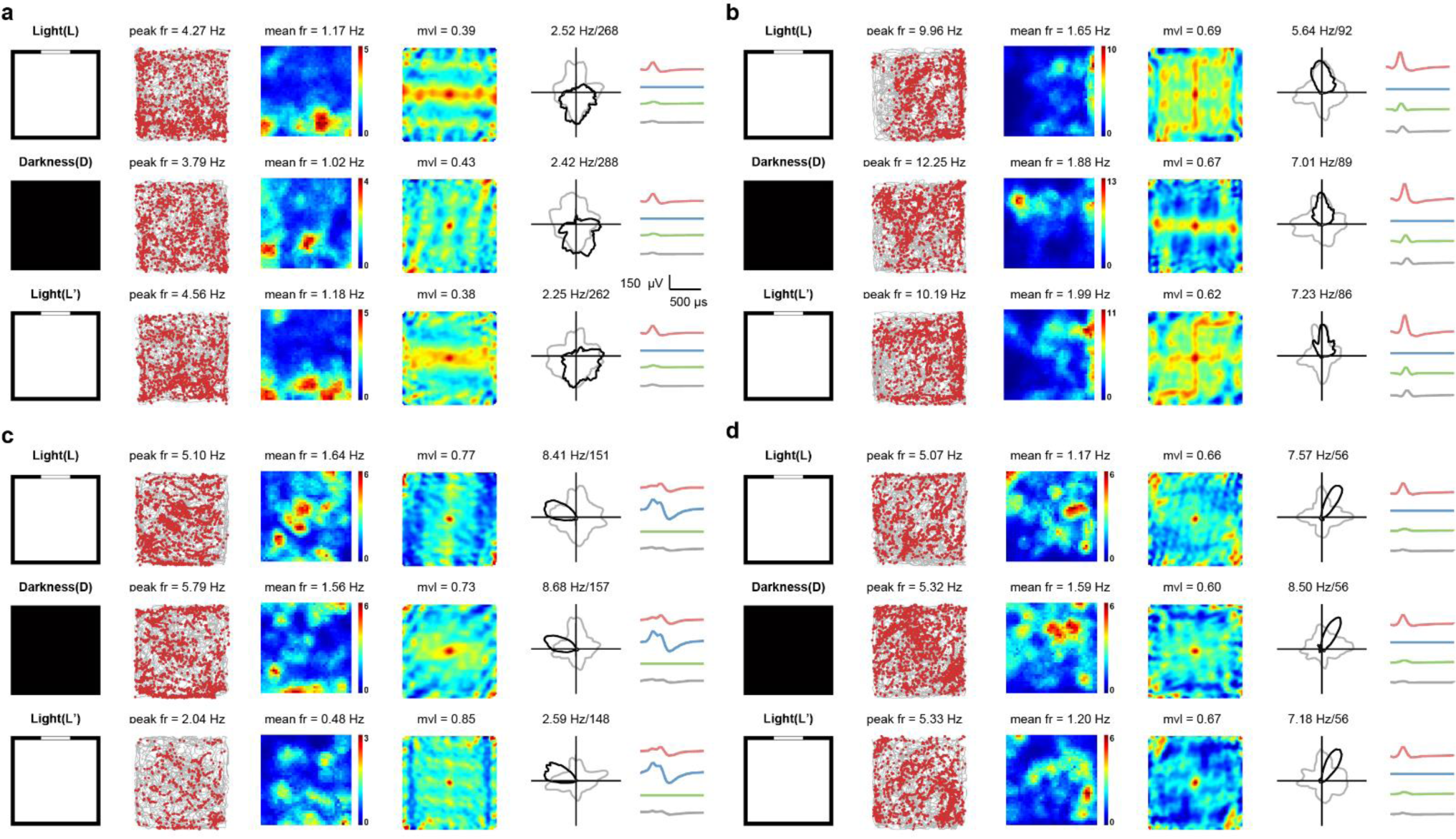
Stable spatial tuning properties of V2 head-direction cells in the darkness. **a-d**, Responses of four representative V2 head-direction cells in the L-D-L’ conditions. Symbols and notations are similar as before.

**Fig. S27.**
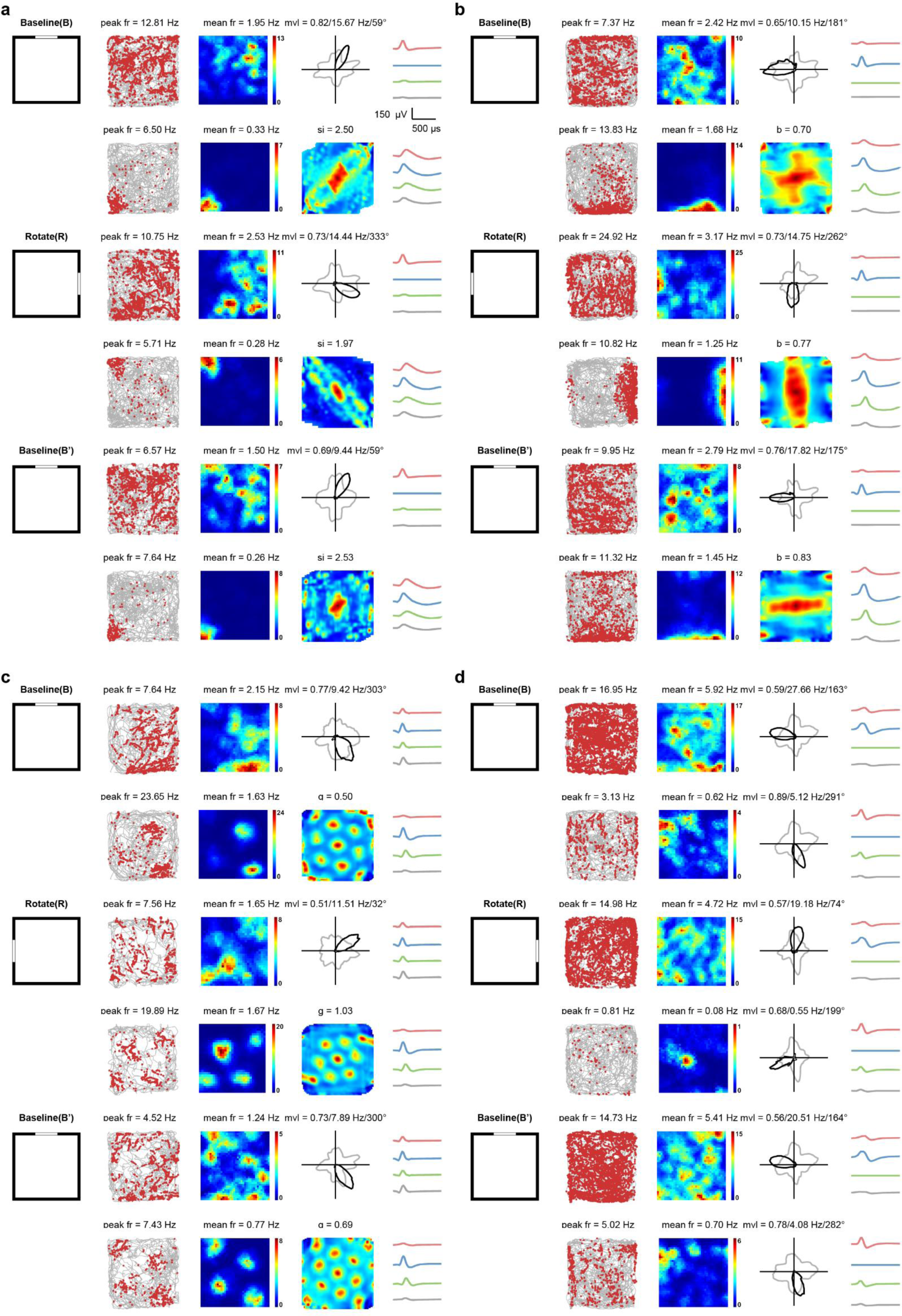
Visual cue control of V2 head-direction cells together with simultaneously recorded other spatial cells. **a**, One V2 head-direction cell with one simultaneously recorded V2 place cell in response to visual landmark manipulation. **b**, One V2 head-direction cell with one simultaneously V2 border cell in response to visual cue rotation. **c**, One V2 head-direction cell with one simultaneously recorded V2 grid cell in response to visual landmark manipulation. **d**, Two simultaneously recorded V2 head-direction cells in response to visual landmark rotation. Symbols and notations are similar as before.

**Fig. S28.**
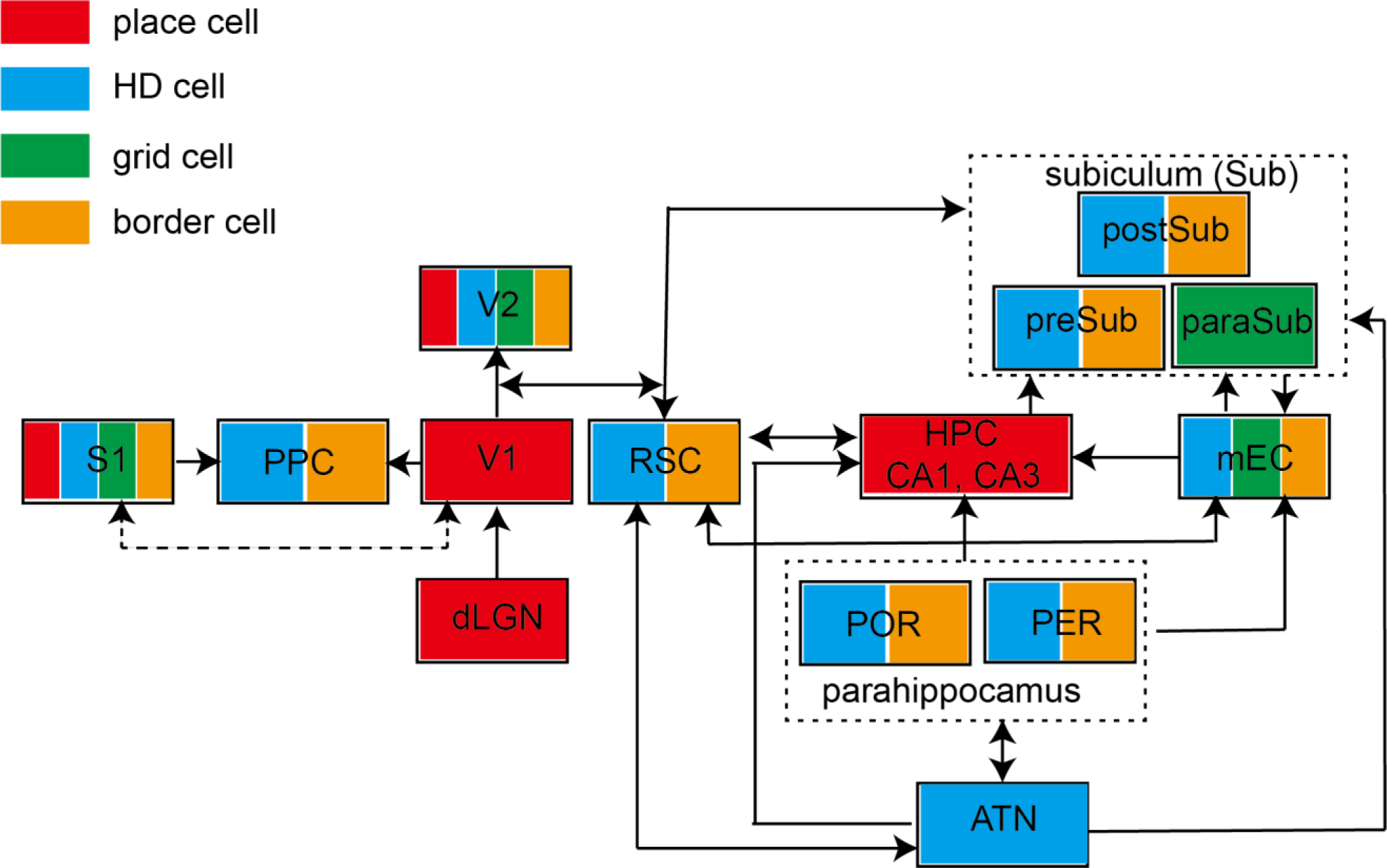
Schematic of identified brain structures with four major types of spatial tunings. Abbreviations: ATN (anterior thalamic nuclei), S1 (primary somatosensory cortex), PPC (posterior parietal cortex), V1 (primary visual cortex), V2 (secondary visual cortex), RSC (retrosplenial cortex), HPC (hippocampus), mEC (medial entorhinal cortex), POR (postrhinal cortex), PER (perirhinal cortex), preSub (presubciculum), paraSub (parasubiculum), postSub (postsubiculum). Arrow indicates the connectivity.

